# West Nile virus response to mosquito and avian biodiversity in rural environments

**DOI:** 10.64898/2026.06.26.734801

**Authors:** Lara Marcolin, Niccolò Ceci, Federica Gobbo, Fabrizio Montarsi, Giulia Chiarello, Ilaria Dorigatti, Moreno Di Marco

**Affiliations:** Department of Biology and Biotechnologies “Charles Darwin”, Sapienza University of Rome, Italy; Istituto Zooprofilattico Sperimentale delle Venezie, Viale dell’Università 10, 35020 Legnaro, Italy; Medical Research Council Centre for Global Infectious Disease Analysis, School of Public Health, Imperial College London, London, United Kingdom

## Abstract

**Context:** The relationship between biodiversity and zoonotic disease risk is a central topic in community ecology, yet empirical evidence in Europe remains scarce and often contradictory compared to North American studies. Addressing this gap is fundamental to better anticipate zoonotic disease dynamics.

**Objectives:** We investigated the transmission dynamics of West Nile virus (WNV) in Veneto (Italy), a major European hotspot. Because this vector-borne pathogen is primarily transmitted by *Culex* mosquitoes and maintained by several avian hosts, we analysed how multiple facets of both avian and mosquito biodiversity influence its transmission.

**Methods:** Using Generalized Additive Models (GAMs) trained on longitudinal entomological and ornithological surveillance data, we modelled the probability of WNV presence in mosquito pools as a function of host and vector community structure. To isolate the effects of biodiversity, we explicitly controlled for climatic and landscape covariates.

**Results:** In agricultural landscapes, we found that higher avian diversity leads to higher viral presence, driven by the dominance of highly competent synanthropic hosts. Conversely, a dilution effect emerges across the broader regional landscape where areas of higher ecological integrity allow for more complex and functionally diverse avian communities. Furthermore, we identified significant vector-mediated regulation, where high abundances of mammophilic vectors effectively suppress viral prevalence through larval competition.

**Conclusions:** Our findings suggest that the dilution effect is a property of intact ecosystems which can be lost, or even locally reversed, in anthropogenically altered environments. Because such habitat degradation fundamentally alters zoonotic transmission dynamics, landscape planning must prioritize ecological restoration. Ultimately, embedding these practices into One Health strategies represents a proactive approach to mitigating disease emergence.

## INTRODUCTION

The relationship between biodiversity and zoonotic disease risk remains one of the fundamental debates in community ecology. Within vector-borne systems, this interaction is often framed through the “dilution effect” hypothesis (Ostfeld & Keesing, 2000), which posits that high species richness suppresses pathogen transmission by decreasing the frequency with which vectors interact with highly competent species (Ostfeld & Keesing, 2012; Rohr et al., 2020). However, empirical evidence remains fragmented and often contradictory (Halliday et al., 2020; Keesing & Ostfeld, 2021). This inconsistency has fuelled a polarized debate with the “amplification effect” hypothesis (Keesing et al., 2006) maintaining that higher diversity can increase prevalence, particularly when newly introduced species are highly competent hosts. Such discrepancies might indicate that the biodiversity–disease relationship is not universal, but rather context-dependent and scale-dependent (Ostfeld & LoGiudice, 2003).

Transmission efficiency in zoonotic vector-borne pathogens relies on the multidimensional interplay between host and vector community structures (Johnson & Thieltges, 2010; LoGiudice et al., 2003; Ostfeld & Keesing, 2012). A critical, yet poorly understood, driver of this context-dependency is anthropogenic disturbance, which acts as a selective filter on host communities (Halliday et al., 2020; Ferraguti et al., 2023). In modified landscapes, disturbance leads to biotic homogenization, favouring widespread, generalist host species (Gibb et al., 2020; Newbold et al., 2018). This occurs because anthropogenic pressures select for synanthropic generalists with life-history traits that frequently correlate with high host competence, such as fast reproduction (Gibb et al., 2020; Johnson et al., 2012; Previtali et al., 2012). Anthropogenic habitat degradation also reshapes vector communities, again favouring a few dominant, and highly competent species (Ferraguti et al., 2016; Perrin et al., 2022). This ecological dominance might erode vector community evenness and focus transmission within a narrow group of efficient species, thereby increasing the probability of pathogen circulation (Hermanns et al., 2023; Martínez-de La Puente et al., 2018). Consequently, characterizing disease risk requires moving beyond taxonomic counts to integrate functional traits and ecological integrity into the analysis of multi-trophic transmission cycles (Di Marco et al., 2026).

West Nile virus (WNV) ecology provides a perfect example of a complex transmission cycle. Primarily transmitted by *Culex* vectors and maintained by several avian hosts, WNV infects humans and horses as dead-end hosts (Keesing & Ostfeld, 2021; Kilpatrick et al., 2005). Evidence of a dilution effect from high bird diversity has been reported for WNV in North America (Allan et al., 2009; Ezenwa et al., 2006; Swaddle & Calos, 2008), but results in Europe are scarce and often contradictory (Ferraguti et al., 2021; Wang et al., 2025). While various studies have integrated host and vector metrics (Adelman et al., 2022; Ferraguti et al., 2021), the functional synergy between these groups across gradients of anthropogenic disturbance remains largely unexplored. Moreover, the relative importance of host versus vector communities in driving WNV circulation is still largely unknown. Recent findings in North America suggest that structural shifts in vector communities—such as the turnover between primary vectors like *Cx. pipiens* and *Cx. tarsalis* driven by land-use changes—can be significantly more predictive of zoonotic risk than avian assemblages, with bird diversity often showing negligible influence on infection rates (Adelman et al., 2022).

In Europe, the Mediterranean basin is currently a primary area for WNV circulation, with Italy as one of the major hotspots in terms of human outbreaks (Rizzo et al., 2016). Within this context, the Veneto region represents a critical risk area due to a mosaic of agricultural matrices and extensive wetlands which sustain high vector and host densities (Giesen et al., 2023; Marini et al., 2018; Riccardo et al., 2022). Yet, despite recurrent WNV outbreaks since the late 2000s, it is still unclear how local biodiversity shapes viral activity across this region. In this study, we explore how several biodiversity facets modulate WNV presence across a gradient of anthropogenic disturbance in Veneto. We investigate the probability of viral presence in mosquito pools as a function of host and vector community structure. By integrating field-collected longitudinal data with theoretical species richness models, we test the following hypotheses:

*H1. The biodiversity–disease relationship is a scale-dependent outcome of anthropogenic disturbance. a) We hypothesize that in agricultural landscapes higher avian biodiversity leads to viral amplification due to the dominance of synanthropic hosts (Gibb et al., 2020). b) Conversely, a dilution effect emerges across broader ecological gradients spanning high environmental integrity, which allows for more complex community structure (Marcolin et al., 2024)*.

*H2. Vector-mediated regulation. a) We hypothesize that transmission in agricultural landscapes is regulated primarily by vector community assembly (Adelman et al., 2022), given the limited overall variability of host communities. b) We hypothesize that a varied composition of vector communities, especially when there is a high presence of mammophilic vectors, reduces WNV circulation through a combination of interspecific larval competition and an increased proportion of bites wasted on dead-end hosts by mammophilic adults (Carrieri et al., 2003; Costanzo et al., 2005)*.

## METHODS

### Biological predictors of WNV presence

We investigated the relationship between biodiversity and WNV circulation in Veneto (Italy) by integrating entomological, ornithological, and environmental data using mosquito trap stations as reference spatial units (Fig. 1).

**Figure 1.**
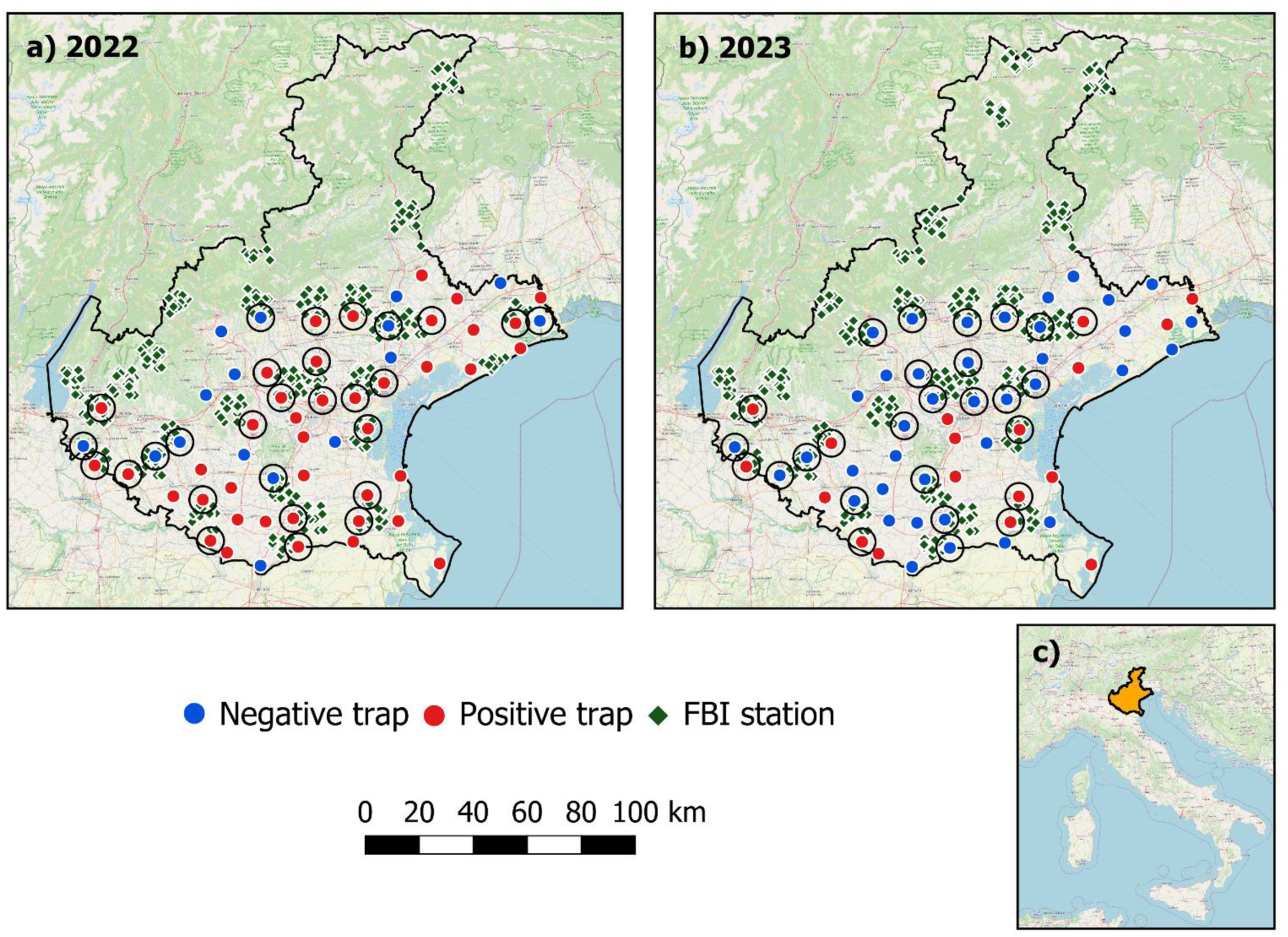
Entomological and ornithological data for (a) 2022 and (b) 2023. Mosquito trap stations are shown in red (WNV-positive) and blue (WNV-negative), and Farmland Bird Index (FBI) survey stations in green. Circular buffers (5 km radius) indicate the traps included in the study and the spatial unit used to integrate entomological and bird data. The inset map (c) shows the location of the study area within the national context of Italy.

### Viral and biodiversity data

We obtained entomological data from the Regional West Nile virus and Usutu virus (USUV) surveillance programme in the Veneto region during the 2022 and 2023 transmission seasons (Gobbo et al., 2025). This monitoring followed the Italian national plan for the prevention, surveillance and response to arboviral diseases 2020–2025 (Ministero della Salute, 2019). Mosquitoes were sampled bi-weekly from mid-May to mid-October using a network of CDC-CO₂ and gravid traps. Female specimens of the *Cx. pipiens* complex, *Aedes albopictus*, and *Ae. caspius* were pooled by species (up to 100 individuals) and screened for orthoflaviviruses using a one-step SYBR green-based real-time RT-PCR assay. Comprehensive technical protocols regarding morphological identification and laboratory procedures are detailed in Appendix S1: Section S1.

Ornithological data for 2022 and 2023 were obtained from the national Farmland Bird Index programme, coordinated by the Rete Rurale Nazionale in collaboration with LIPU (Rete Rurale Nazionale & LIPU, 2023). Avian surveys were conducted only in agricultural landscapes, following the unlimited-distance point count methodology (Blondel et al., 1981). Within each 10×10 km UTM cell, 15 point count stations were positioned in 1×1 km cells selected through spatial randomisation (Fornasari et al., 2002). Surveys were conducted during the breeding season, with timing adjusted for latitude and elevation (see Appendix S1: Section S2). As bird surveys were conducted independently from entomological surveys and limited to agricultural areas, we employed circular buffers of 5 km and 10 km (as a sensitivity analysis) around each entomological trap to associate these to bird data. In total, we had 55 associations at the 5 km scale (28 in 2022, 27 in 2023; Figure 1), and 89 associations at the 10 km scale (47 in 2022, 42 in 2023).

Based on the associations between entomological data and bird survey data, we established two primary spatial settings: (i) an “agricultural” setting, limited to entomological traps associated with at least one bird survey location within the agricultural landscape; (ii) a “full” setting, focussed on the entire entomological dataset defined using all available entomological trap stations monitored over two years (2022–2023). In fact, while we could derive fine-scale bird biodiversity metrics only for the agricultural landscape, we could analyse vector biodiversity at both scales. Moreover, by looking at the broader landscape, we were able to extend our analysis across a significantly wider ecological gradient, spanning both modified environments and semi-natural systems. To characterise the environmental context of these two spatial settings, we used the Biodiversity Intactness Index (BII; Newbold et al., 2016) as a proxy for ecological degradation (Appendix S1: Section S3 and Figure S1).

### Estimating biodiversity metrics

To identify which aspects of avian biodiversity may modulate WNV transmission, we derived a suite of metrics designed to capture different dimensions of host community structure. Based on bird survey data associated with each entomological trap, we calculated relative empirical avian species abundances from point count data, assuming that standardized counts reflect local community composition. We calculated taxonomic diversity, including species richness (*S_b_*), Shannon diversity (*H_b_*), Simpson diversity (1 - *D_b_*), and evenness (*J_b_*) to evaluate how the numerical distribution and variety of species influence transmission potential. These metrics allow us to assess whether viral risk is modulated by the simple presence of species or by the relative dominance of certain individuals within the community. Beyond taxonomic counts, we quantified avian functional richness to represent the variety of ecological roles and life-history strategies present in the community (Legras et al., 2018). This metric is critical for understanding how functional traits, such as reproductive investment and dispersal ability, correlate with host competence and influence the dilution or amplification of the virus (Marcolin et al., 2026). Functional richness was calculated by mapping local species onto a multidimensional functional space (Carmona et al., 2024), defined via Principal Component Analysis ordination using a matrix of 24 avian functional traits (Appendix S1: Table S1; Mammola et al., 2021). Detailed information regarding the mathematical formulations of biodiversity metrics and trait imputation is provided in Appendix S1: Sections S4 and S5. To address the scale-dependency of the dilution effect, we incorporated “potential” avian species richness derived from published species distribution models (Si-moussi & Thuiller, 2024). This metric allowed us to explore WNV dynamics across a broader gradient of ecological disturbance extending beyond the degraded rural-agricultural matrix captured by empirical field surveys.

To capture the local entomological context and its influence on WNV circulation, we derived several metrics related to vector abundance and community structure from the raw mosquito count data described in Appendix S1: Text S4. We calculated total abundance as the number of mosquitoes captured per trap per night, aggregated across the May–October season, to serve as a proxy for overall vector population density. To evaluate the specific roles of primary and secondary vectors, we determined the relative abundance of four species critical to local transmission (Gobbo et al., 2025): *Cx. pipiens*, *Cx. modestus*, *Aedes albopictus*, and *Ae. caspius*. This metric was calculated as the count of a specific species divided by the total number of all mosquitoes captured in the same trap and year. This allowed us to assess how WNV presence is modulated by the dominance of avian-feeding vectors (*Cx. pipiens*, *Cx. modestus*) versus mammophilic bridge vectors (*Aedes sp.*) in the enzootic cycle (Muñoz et al., 2011; Soto & Delang, 2023; Vinogradova, 2000). Beyond simple counts, we integrated mosquito species richness (*S_m_*) and community evenness (*J_m_*) to investigate the vector-mediated dilution hypothesis. These community metrics provide insight into the multi-trophic regulation of transmission; for instance, high evenness or the presence of non-primary vectors may indicate competitive interactions (Carrieri et al., 2003; Costanzo et al., 2005). By analyzing these facets of the entomological community, we can decouple the influence of vector diversity from absolute abundance and test whether higher community complexity effectively suppresses viral prevalence.

### Environmental predictors of WNV presence

WNV transmission is governed by the complex interplay between host community structure and environmental conditions (Vargas Campos et al., 2025). To robustly isolate the effects of biodiversity, we controlled for climatic and landscape predictors known to regulate mosquito life cycles, viral replication, and host–vector contact rates (Brown et al., 2014; Esser et al., 2019; Giesen et al., 2023; Perrin et al., 2022). Climatic data, comprising seasonal mean temperature, minimum relative humidity (averaged across the season), and cumulative precipitation, were obtained from the ARPA Veneto meteorological network (https://www.arpa.veneto.it) and averaged across the *Cx. pipiens* activity season (May–October) for 2022 and 2023.

Landscape characteristics are also known to regulate WNV circulation, affecting both vector and host ecology by determining the availability of breeding sites for mosquitoes (Giesen et al., 2023; Perrin et al., 2022) and the distribution of foraging and nesting resources for birds (Rigal et al., 2023). We retrieved land-cover data from the ESA WorldCover 2021 dataset (Zanaga et al., 2022) to calculate the proportions of seven land cover classes, including critical wetlands and irrigated croplands at a 10 m spatial resolution. Vegetation productivity, which sustains high *Culex* densities (Brugueras et al., 2020; Esser et al., 2019), was represented through the seasonal mean Normalized Difference Vegetation Index (NDVI) derived from Landsat imagery (Hemati et al., 2021). Anthropogenic pressure and altered hydrology were quantified using the Copernicus Imperviousness Density layer (Copernicus Land Monitoring Service, 2018), as soil sealing is a primary driver of *Cx. pipiens* suitability (Gangoso et al., 2020; Marcolin et al., 2025). To align with the spatial scales used for the avian biodiversity metrics, these proportions were extracted within both 5 km and 10 km circular buffers around each entomological trap.

### Modelling WNV presence

We ran Generalized Additive Models (GAMs) to assess the relationship between biodiversity metrics and the probability of detecting WNV-positive mosquitoes, while controlling for environmental covariates. Models were fitted using the *mgcv* package in R (Wood, 2000), with a binomial error distribution and logit link function. Predictor smooth terms were estimated using thin plate regression splines, with smoothing parameters selected via restricted maximum likelihood (REML). Continuous predictors were centred and scaled prior to analysis. The response variable was binary, representing the presence (1) or absence (0) of WNV in mosquito pools sampled at each trap station and for each year. A trap was considered WNV-positive if it contained at least one positive mosquito pool, regardless of the species. To reduce possible overfitting and ensure that modelled relationships remained biologically interpretable, we constrained the smoothness parameter to *k*=3, roughly corresponding to quadratic effects. We accounted for spatial autocorrelation among sampling locations by including a tensor product smooth of easting and northing coordinates as a trend surface (Fletcher & Fortin, 2018).

To isolate the specific contributions of biodiversity, we first developed a parsimonious environmental base model. We addressed multicollinearity by excluding variables with pairwise correlations > 0.7 (Appendix S1: Figure S3), resulting in a set of candidate predictors comprising humidity, temperature, NDVI, cropland, imperviousness, sparse vegetation, and wetlands. Following a saturated model fit, we applied a stepwise backward elimination, retaining only predictors with strong explanatory power (p < 0.05).

We extended the environmental base model through a multi-step procedure to test the individual and joint roles of biological variables. To evaluate the role of vector biodiversity, we incorporated one of six mosquito biodiversity indicators at a time, repeating the models on the full landscape (n=114 entries) and the agricultural landscape (n=55 entries). Similarly, we assessed the contribution of avian biodiversity to WNV presence in the agricultural landscape, where Farmland Bird Index monitoring stations were placed. We accounted for sampling effort to control for potential biases arising from the varying number of bird monitoring stations associated with each entomological station. Sampling effort was estimated as the number of bird survey stations associated with each entomological trap and was included as a covariate when fitting the models. To ensure that our findings were not artifacts of sampling bias, we also performed a sensitivity analysis by focussing only on sites with the highest bird data completeness. In this case, we estimated sample completeness across all bird survey stations associated with each entomological trap using the Chao2 estimator (Chao & Jost, 2012). Conversely, potential species richness, which derives from consistent modelling estimates rather than direct counts, was considered independent of local sampling effort and could be tested in both the agricultural and full landscapes, to capture a broader ecological integrity gradient.

Finally, we investigated the integrated effects of host and vector communities by fitting joint models. Given its role as the primary enzootic amplifier in the region (Mancini, 2017), the relative abundance of *Cx. pipiens* was selected *a priori* to represent the vector component and combined with each of the six avian biodiversity variables. This approach allowed us to effectively decouple the influence of host community structure from the confounding effect of primary vector pressure. Partial dependence plots (*visreg* package; Breheny and Burchett, 2017) were used to visualize the relationships between biodiversity predictors and WNV presence, while discounting the effects of environmental factors and sampling size (for the agricultural model).

The predictive performance of all models was evaluated using a Leave-One-Out Cross-Validation (LOOCV) framework, ensuring an unbiased estimate of out-of-sample error by utilizing the maximum available data for training. For each dataset of *n* units, the model was iteratively trained on *n*-1 observations and tested on the omitted unit. Global performance was quantified using the True Skill Statistic (TSS; Allouche et al., 2006) with a standard binarization threshold of 0.5.

## RESULTS

The spatial sensitivity analysis confirmed the 5 km buffer as the optimal scale for extracting landscape predictors (Appendix S1: Tables S2–S4; Appendix S1: Figures S4-S5). Minimum adequate environmental models explained 33.7% of the deviance for the agricultural landscape dataset and 23.3% for the full dataset, both identifying a significantly higher probability of WNV presence in 2023 compared to 2022 (p < 0.001). In the agricultural landscape dataset, imperviousness (χ² = 7.14, p = 0.011) and sparse vegetation (χ² = 4.22, p = 0.025) emerged as key predictors of mosquito positivity. Across the broader regional gradient, cropland, sparse vegetation, and wetland proportions were identified as significant environmental predictors of mosquito positivity (Appendix S1: Table S4).

### Biodiversity–disease relationship and anthropogenic disturbance

Within the agricultural landscape, models incorporating avian biodiversity metrics outperformed the base environmental model, achieving moderate predictive performance with 35.5%-54.6% deviance explained (Table 1; Appendix S1: Table S5). Models’ evaluation similarly led to moderately good outcomes in terms of TSS (0.27-0.48; Appendix S1: Table S6). The inclusion of observed species richness led to the best performing model, with much higher predictive power compared to potential species richness (54.6% *vs* 35.5%) indicating a better ability to capture local biodiversity levels rather than modelled distributions.

**Table 1.** Summary results of GAMs predicting the presence of WNV in mosquito pools based on a set of environmental characteristics (i.e. “base model” in the text) and an individual avian biodiversity metric. All models were fitted using the agricultural landscape dataset.

| Biodiversity variable | Chi.sq | p-value | R<sup>2</sup> (adj.) | Deviance explained |
| --- | --- | --- | --- | --- |
| Avian species richness | 10.843 | 0.002 | 0.558 | 54.6% |
| Avian functional richness | 8.526 | 0.006 | 0.393 | 38.5% |
| Avian Shannon index | 7.681 | 0.009 | 0.493 | 50.8% |
| Avian Simpson index | 1.876 | 0.107 | 0.386 | 40.2% |
| Avian evenness | 1.832 | 0.104 | 0.388 | 40.4% |
| Potential avian species richness | 1.241 | 0.137 | 0.358 | 35.5% |

Within the agricultural landscape, the partial dependence plots of all observed avian biodiversity indicators revealed non-linear positive associations with WNV presence (Figure 2). In particular, WNV risk increased rapidly from low to intermediate values of species richness until reaching a high plateau. Functional richness and the Shannon index followed a more gradual, non-linear increase toward saturation. The Simpson index and avian evenness exhibited a steady, near-linear positive association. Collectively, these trends indicate a consistent amplification effect across all facets of avian biodiversity within the rural-agricultural matrix, consistent with our Hypothesis 1a. The sensitivity analysis performed across different levels of sample completeness confirmed that these observed trends are qualitatively stable (Appendix S1: Figures S6-S7), with richness and diversity metrics remaining positively associated with WNV presence. Avian evenness proved to be the only exception, showing higher sensitivity to sampling effort; consequently, its results must be interpreted with caution.

**Figure 2.**
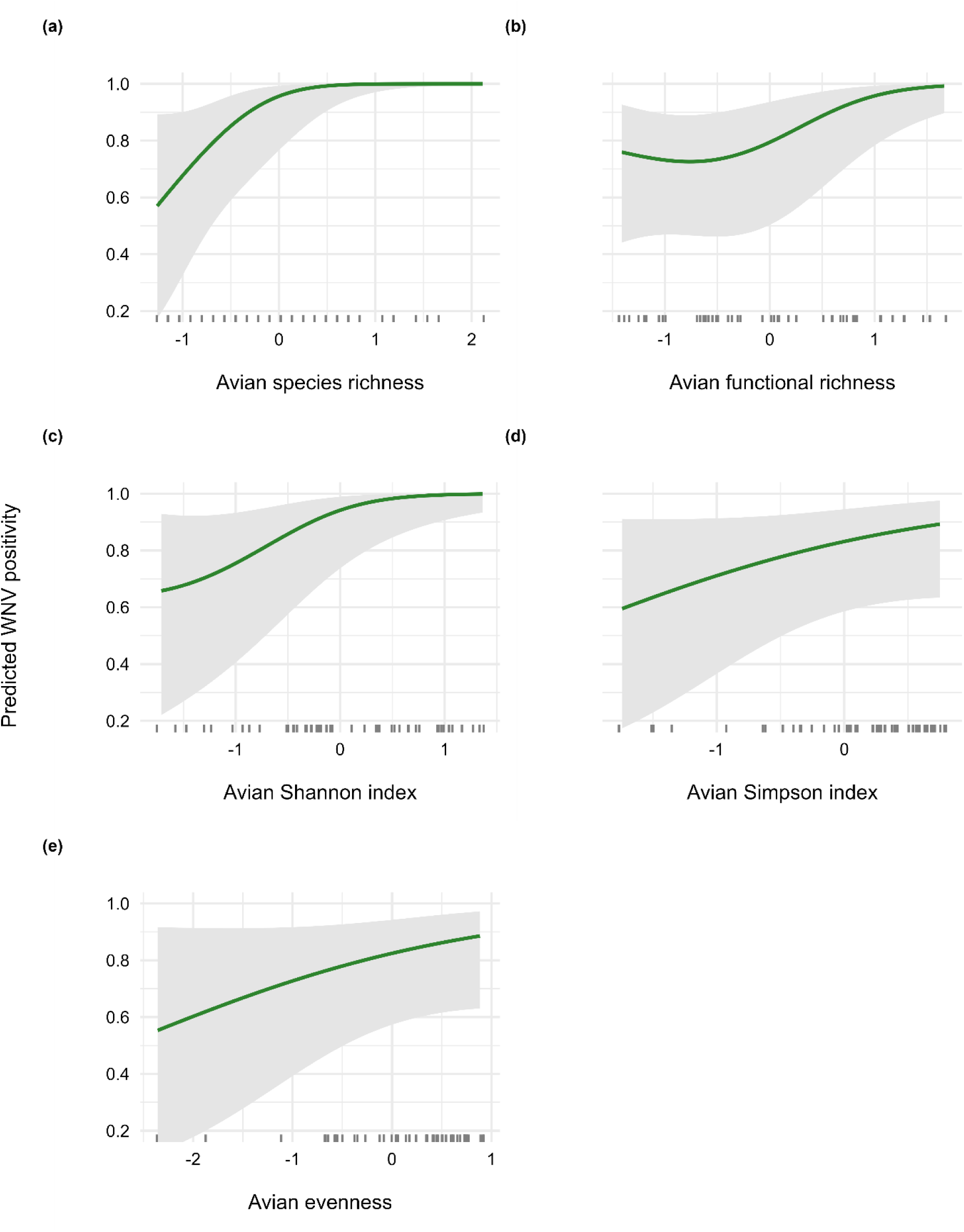
Predicted probability of WNV positivity in mosquitoes as a response to different avian biodiversity metrics. Curves are plotted over the central 95% of observed values for each metric (observation densities are reported at the bottom of each plot). Shaded grey areas indicate 95% confidence intervals. Values on the x axis have been rescaled for analysis. These results are derived from avian biodiversity models fitted on the agricultural landscape dataset.

Within the broader landscape, potential species richness had slightly lower predictive performance than in the agricultural landscape (29.7% deviance explained *vs* 35.5%; Appendix S1: Tables S5 and S7) but much higher evaluation results in terms of TSS (0.54 *vs* 0.34; Appendix S1: Tables S6 and S8). At this scale, potential richness was also a significant predictor of WNV presence (p<0.01; Appendix S1: Table S7), unlike the agricultural landscape (Table 1). The influence of potential avian richness on WNV presence also shifted when looking at the broader landscape rather than the agricultural landscape (Figure 3; Appendix S1: Figure S8). While the relationship was weak within the restricted agricultural matrix, a strong negative association emerged when looking at the full landscape. This result confirms the scale-and context-dependency of the biodiversity-disease association (Hypothesis 1b).

**Figure 3.**
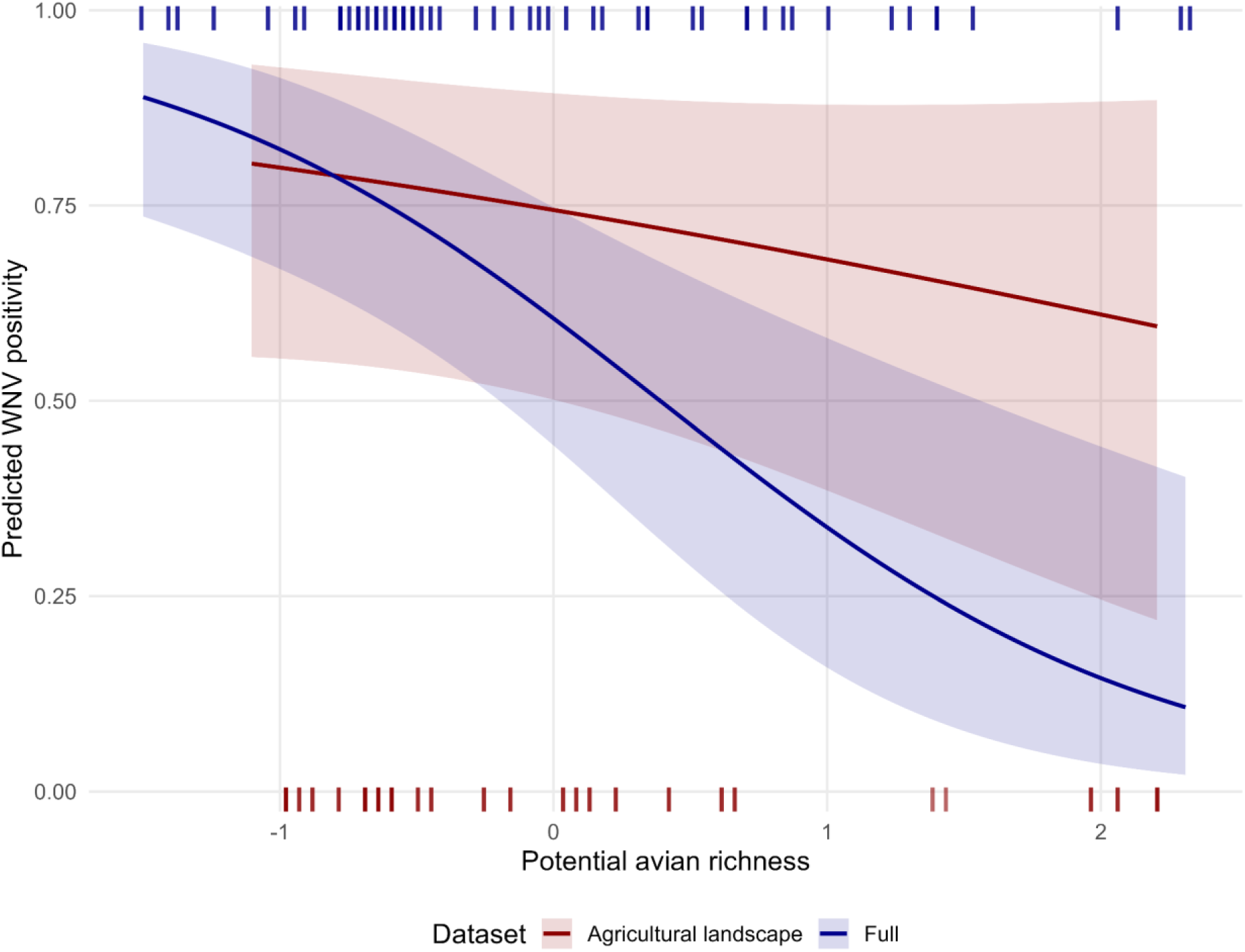
Predicted probability of WNV positivity in mosquitoes as a response to potential avian species, using the agricultural landscape dataset and (red) the full dataset (blue). Curves are plotted over the central 95% of observed values for each metric (observation densities are reported at the bottom of each plot). Shaded areas indicate 95% confidence intervals. Values on the x axis have been rescaled for analysis.

### Vector-mediated regulation

Incorporating mosquito biodiversity metrics generally improved the performance of the base models in the agricultural landscape (deviance explained 35.7%-43.4%), but was less influential in the full landscape, which contrasts with Hypothesis 2a (23.3%-31.8%; Table 2; Appendix S1: Tables S9-S10). Models including *Ae. albopictus* and *Oc. caspius* abundance achieved the best validation performance with a TSS of 0.35 and 0.41, respectively (Appendix S1: Table S11).

**Table 2.**
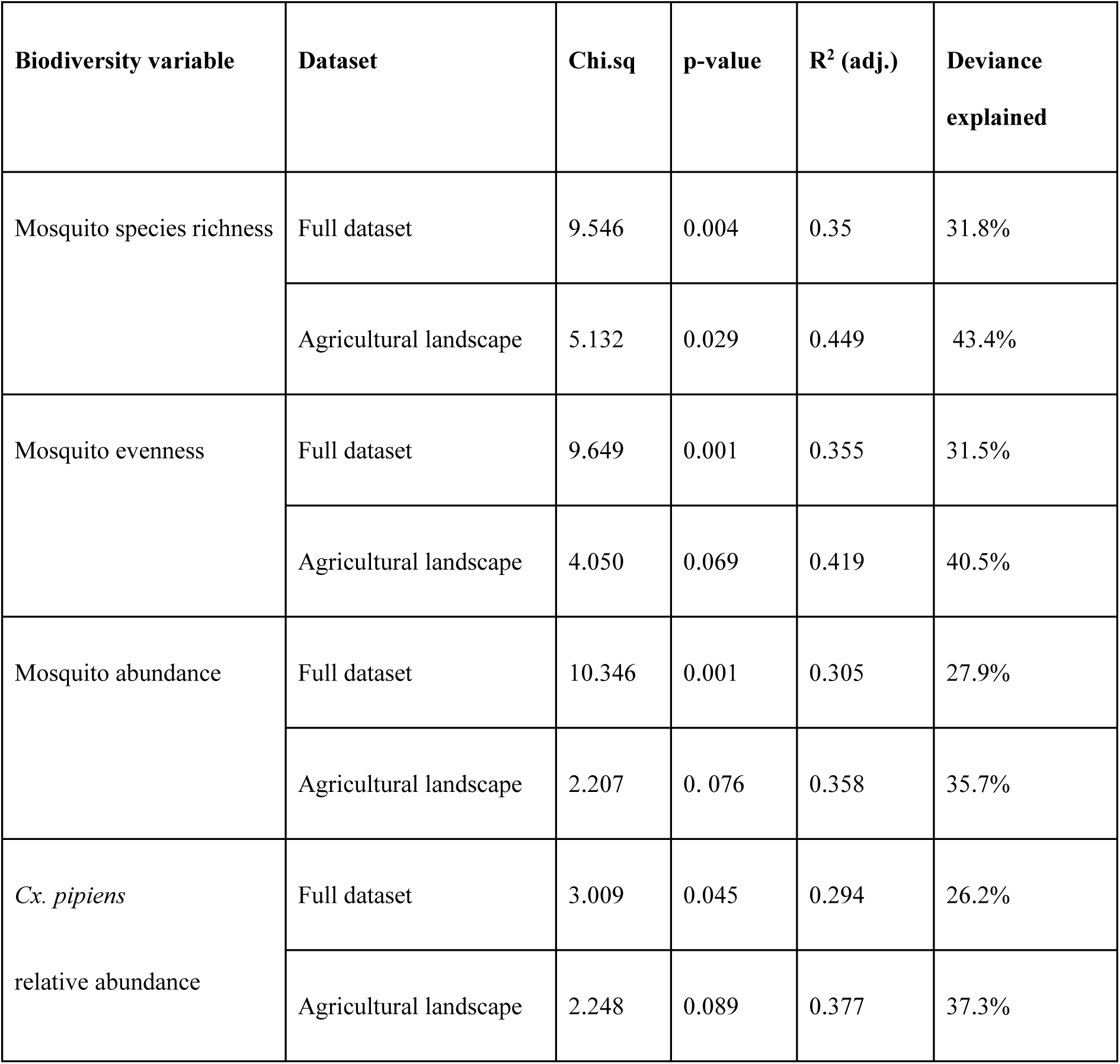
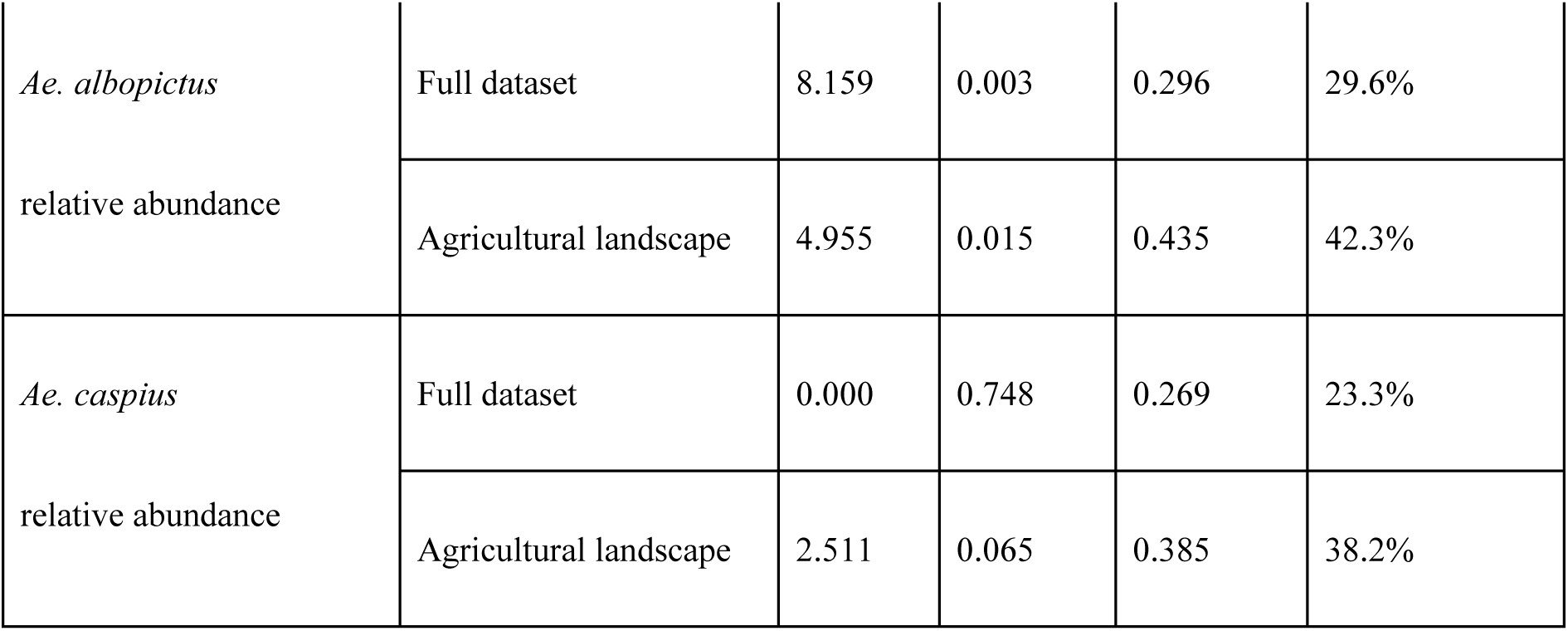
Summary results of GAMs predicting the presence of WNV in mosquito pools based on a set of environmental characteristics (i.e. “base model” in the text) and a given mosquito biodiversity metric. Several biodiversity metrics were tested, one at a time. Results are presented for both the full dataset and the agricultural landscape dataset (referring to traps where both mosquito and bird data were available).

Partial dependence plots revealed distinct response patterns of WNV presence to mosquito community metrics (Figure 4; Appendix S1: Figure S9), in particular the unimodal response of WNV presence to mosquito richness. Conversely, mosquito evenness and *Ae. albopictus* abundance exhibited strong, monotonic negative associations with WNV presence, supporting Hypothesis 2b on vector-mediated modulation of transmission.

**Figure 4.**
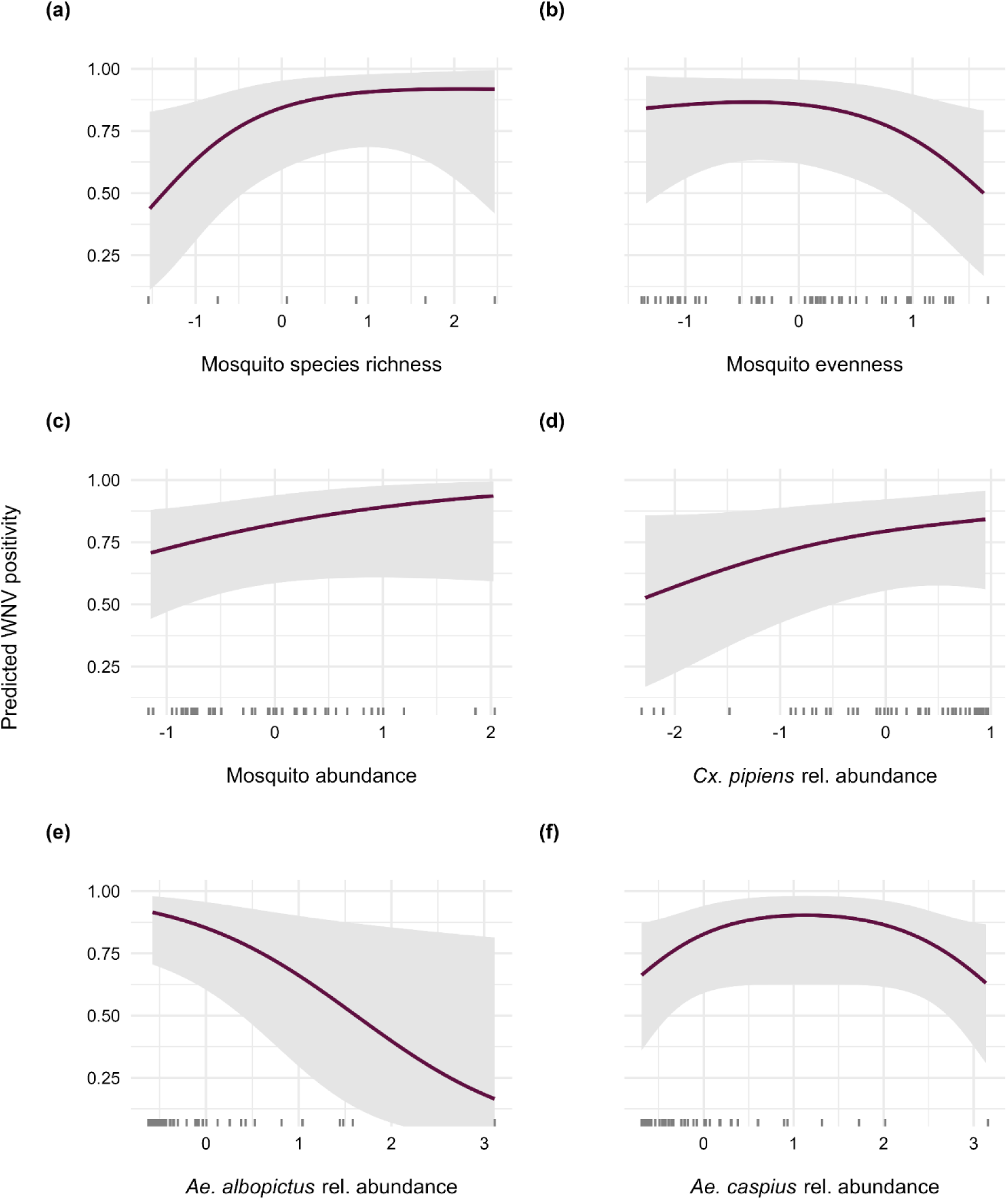
Predicted probability of WNV positivity in mosquitoes as a response to different mosquito biodiversity metrics. Curves are plotted over the central 95% of observed values for each metric (observation densities are reported at the bottom of each plot). Shaded grey areas indicate 95% confidence intervals. Values on the x axis have been rescaled for analysis. These results are derived from entomological models fitted on the agricultural landscape dataset.

Integrated host-vector models in the agricultural landscape dataset showed higher relevance of avian biodiversity metrics (especially species richness) compared to mosquito metrics (including *Cx. pipiens a*bundance), with a deviance explained of 54.6% *vs* 37.3% (Table 3; Appendix S1: Table S12) and TSS 0.49 *vs* 0.35 (Appendix S1: Table S13). Again, this contrasts with our original Hypothesis 2a.

**Table 3.** Summary results of GAMs predicting the presence of WNV in mosquito pools based on a set of environmental characteristics (i.e. “base model” in the text), mosquito abundance, and individual avian biodiversity metrics. All models were fitted using the agricultural landscape dataset.

| **Biodiversity variable** | **Chi.sq** | **p-value** | **R<sup>2</sup> (adj.)** | **Deviance explained** |
| --- | --- | --- | --- | --- |
| *Cx. pipiens* abundance +<br>Avian species richness | 10.843 | 0.002 | 0.558 | 54.6% |
| *Cx. pipiens* abundance +<br>Avian functional richness | 8.329 | 0.007 | 0.397 | 39.4% |
| *Cx. pipiens* abundance +<br>Avian Shannon index | 7.681 | 0.009 | 0.493 | 50.8% |
| *Cx. pipiens* abundance +<br>Avian Simpson index | 1.653 | 0.128 | 0.409 | 42% |
| *Cx. pipiens* abundance +<br>Avian evenness | 1.680 | 0.117 | 0.415 | 42.5% |
| *Cx. pipiens* abundance +<br>Potential avian species richness | 0.751 | 0.186 | 0.391 | 38.5% |

We verified the composition of mosquito communities, especially the abundance of mammophilic *Ae. albopictus* and *Oc. caspius,* were stronger predictors of WNV presence in agricultural landscapes compared to overall mosquito abundance or richness, achieving a higher performance both in terms of predictive performance (42.3%-38.2%) and TSS (0.35-0.40), again supporting Hypothesis 2b.

## DISCUSSION

We developed models integrating entomological, ornithological, and environmental data to investigate the relationship between biodiversity and the circulation of WNV in Veneto, a major European hotspot of WNV incidence (Riccardo et al., 2022). Our results indicate that the dilution effect is a property of intact ecosystems that is lost or reversed in anthropogenically altered environments. In these degraded landscapes, viral activity is governed by the interplay between simplified host communities, vector dominance, and land-use change, consistent with broader theory linking biodiversity loss to pathogen amplification in degraded ecosystems (Gibb et al., 2020; Gottdenker et al., 2014; Newbold et al., 2018).

This context-dependency confirms that the biodiversity–disease relationship is a scale-dependent outcome of anthropogenic disturbance, as proposed in our first hypothesis. Within the rural-agricultural matrix, field-derived metrics revealed a significant positive non-linear relationship between WNV positivity and bird diversity (Hypothesis 1a). This local amplification effect reflects the biotic homogenization of degraded systems. National monitoring via the Farmland Bird Index confirms this ecological crisis: while indicators of ecosystem health decreased by over 50% between 2000 and 2023 (Rete Rurale Nazionale & Lipu, 2024), disturbance-tolerant generalists like the Eurasian magpie (*Pica pica*) and hooded crow (*Corvus corone*) have increased in abundance. These synanthropic Passeriformes are highly competent WNV hosts capable of sustaining high viremia and disproportionately driving viral amplification in the region (Gobbo et al., 2025). In contrast, potential avian richness exhibited a consistent negative association with WNV detection across the full regional landscape, representing the regional “dilution potential” (Hypothesis 1b). While this dilutive trend was most robust across the full regional gradient, it persisted, albeit without reaching statistical significance, even within the lower-integrity rural matrix where empirical species richness showed amplification. This discrepancy suggests that the dilution effect is an emergent property of ecological integrity that requires a sufficient breadth of the biodiversity gradient to be empirically detected (Halliday et al., 2020).

In degraded environments, local environmental filters and the dominance of a few highly competent reservoirs likely introduce sufficient stochastic noise to suppress the statistical signal of dilution. While potential avian richness represents the regional species pool that could be expected to persist in a certain environment, field-based surveys capture the realized community that has successfully passed through local environmental filters like intensive agriculture, hunting, human disturbance, and habitat fragmentation. These fine-scale determinants of species persistence are often overlooked by distribution models, which typically rely on coarse environmental predictors (Lembrechts et al., 2019). Such local factors are, however, the primary drivers of bird community structure in highly fragmented landscapes like the rural-agricultural matrix of the Veneto region (Rigal et al., 2023). In these degraded matrices, community assembly is biased toward generalist species with life-history traits that often correlate with high host competence (Gibb et al., 2020; Johnson et al., 2015).

Our findings also demonstrated that mosquito community structure significantly influences the probability of detecting WNV-positive pools, supporting the regulatory mechanisms proposed in our second hypothesis (H2b). While viral circulation closely tracks the relative abundance of *Cx. pipiens*, the principal enzootic vector in Italy (Mancini, 2017), the strong negative associations between WNV detection and both mosquito evenness and *Ae. albopictus* abundance supports a vector-mediated dilution. This pattern likely stems from competitive interactions, such as the larval superiority of *Ae. albopictus* over *Cx. pipiens* under resource-limited conditions (Carrieri et al., 2003; Costanzo et al., 2005). Furthermore, although *Ae. albopictus* is a competent vector for several arboviruses, its strong preference for mammalian hosts limits its potential to amplify WNV compared with *Culex* species (Muñoz et al., 2011; Vogels et al., 2017).

Contrary to our initial expectations (Hypothesis 2a) based on recent findings in sub-urban locations in North America (Adelman et al., 2022), we found that in the rural-agricultural matrix of Veneto, avian biodiversity, particularly species richness, is a far more dominant driver of WNV presence than vector abundance. This suggests that the relative importance of host versus vector communities may vary significantly between geographical regions. This discrepancy is likely explained by the specific environmental gradients investigated. In the system studied by Adelman et al. (2022), the transition from sub-urban to agricultural areas causes a sharp decline in the abundance of *Cx. pipiens*, the primary WNV vector in that region. Because the primary vector’s abundance becomes a limiting factor in those agricultural sites, vector metrics emerge as the strongest predictors of viral activity. In contrast, our study focuses strictly on a rural-agricultural gradient where *Cx. pipiens* remains the ubiquitous and overwhelmingly dominant enzootic vector (Mancini, 2017; Gobbo et al., 2025). We hypothesize that, because primary vector pressure is relatively constant across our study area and might not act as a limiting bottleneck, the local structural variation of the highly competent avian community emerges as the overarching driver of WNV transmission. This pattern is strongly supported by Ferraguti et al. (2021), who found that WNV positive association with avian richness remains unchanged even after incorporating mosquito community metrics into their models.

Our findings have critical implications for WNV surveillance and policy. While large-scale biodiversity conservation and habitat restoration might provide long-term, landscape-level risk reduction, as suggested by the potential richness analysis, the protective effects of biodiversity are largely suppressed in heavily degraded rural landscapes. Here, avian communities are dominated by disturbance-tolerant, highly competent species that drive local viral amplification. Effective management therefore requires a dual strategy: in the short term, targeted mosquito control must be prioritized to control vector populations and reduce transmission in high-risk, human-modified hotspots; in the long term, protecting and restoring diverse avian communities is essential to maintain ecological integrity, allowing the dilution effect to operate across the broader landscape. Collectively, these results highlight the urgent need to embed biodiversity conservation and habitat management directly into public health policies (Caceres-Escobar et al., 2023; Di Marco et al., 2020), moving toward a truly integrated One Health approach.

## FUNDING

The data were collected as part of the Farmland Bird Index project, funded by the Ministry of Agriculture, Food Sovereignty and Forests through the RRN 2014/2022 program, FEASR – European Agricultural Fund for Rural Development. This research was supported by EU funding within the NextGeneration EU-MUR PNRR Extended Partnership initiative on emerging infectious diseases (project no. PE00000007, INF-ACT Spoke4). ID acknowledges funding from the MRC Centre for Global Infectious Disease Analysis (reference MR/X020258/1), funded by the UK Medical Research Council (MRC). This UK funded award is carried out in the frame of the Global Health EDCTP3 Joint Undertaking. ID also acknowledge funding from Wellcome Trust (213494/Z/18/Z, 226072/Z/22/Z, 226727/Z/22/Z and 228185/Z/23/Z). The funders played no role in study design, data collection and analysis, decision to publish, or preparation of the manuscript.

## DATA AVAILABILITY

The datasets generated and/or analysed during the current study along with the essential code employed for statistical modelling are available here: https://doi.org/10.5281/zenodo.18431840.

## COMPETING INTERESTS

The authors declare no conflicts of interest.

## Notes

### Competing Interest Statement

The authors have declared no competing interest.

