## Appendix 1 for "West Nile virus response to mosquito and avian biodiversity in rural environments"

**Appendix S1: West Nile virus response to mosquito and avian biodiversity in rural environments**

*Lara Marcolin[1]*, Niccolò Ceci[1], Federica Gobbo[2], Fabrizio Montarsi[2], Giulia Chiarello[2], Ilaria Dorigatti[3], Moreno Di Marco[1]*

1Department of Biology and Biotechnologies "Charles Darwin", Sapienza University of Rome, Italy

2Istituto Zooprofilattico Sperimentale delle Venezie, Viale dell'Università 10, 35020 Legnaro, Italy

3Medical Research Council Centre for Global Infectious Disease Analysis, School of Public Health, Imperial College London, London, United Kingdom

### Section S1: Entomological sampling and laboratory protocols

Entomological monitoring utilized 57 CDC-$CO_2$-like traps (Italian Mosquito Trap IMT®, PeP, Cantu, Italy) and 7 homemade gravid traps. Captured specimens were transported at 4 °C and morphologically identified under a stereomicroscope using standard taxonomic keys (Severini et al., 2009). All WNV- and/or USUV-positive samples were confirmed by the National Reference Centre for Exotic Diseases of Animals, utilizing a one-step SYBR green-based real-time RT-PCR assay (Ravagnan et al., 2015).

### Section S2: Ornithological sampling

Ornithological data were obtained from the national Farmland Bird Index (FBI) programme, which is coordinated by the Rete Rurale Nazionale in collaboration with LIPU (Rete Rurale Nazionale & LIPU, 2023). Data collection followed the unlimited-distance point count methodology described by Blondel et al. (1981) and adapted for the Italian national breeding bird monitoring scheme (Fornasari et al., 2002). Within each 10×10 km UTM cell, 15 point count stations were positioned in distinct 1×1 km cells selected through a spatial randomisation procedure to ensure unbiased sampling coverage. Each survey consisted of a 10-minute point count conducted during the breeding season, with timing adjusted according to latitude and elevation of sampling stations: generally between 15 May and 30 June, extending to the first week of July for Alpine sites. Counts began shortly after sunrise and were conducted only under favourable weather conditions, avoiding days with strong wind or heavy precipitation. Each station was visited once per year, and observers recorded all birds detected by sight or sound. The final dataset comprised 4,043 individual records from 123 species obtained in 2022 through 415 point counts across 29 UTM cells, and 4.630 records in 2023 from 410 point counts across 34 cells.

### Section S3: Biodiversity Intactness Index

The Biodiversity Intactness Index (BII) quantifies the average abundance of native species relative to an undisturbed state (Newbold et al., 2016). It serves as a proxy for the ecological degradation that reshapes zoonotic communities through biotic homogenization. We used this index to characterize the environmental integrity of the agricultural and full landscape settings, as shown in the distribution curves in Figure S1.

### Section S4: Biodiversity indices

The following standard formulas were used to calculate taxonomic biodiversity metrics for both mosquito and avian communities:

i) Species richness (*S*), calculated as the total number of avian species recorded within each buffer.

ii) Shannon diversity (*H*), which represents how evenly individuals are distributed among different species (Shannon, 1948), measured as follows:

$$H = -\sum_{i=1}^{S} \quad P_i \, ln \, P_i \qquad (1)$$

a. where $P_i$ is the relative abundance of avian species $i$. A larger value of $H$ suggests higher diversity.

iii) Simpson diversity (1-D), which measures the probability that two randomly selected individuals belong to different species (Simpson, 1949) and is measured as follows:

$$1 - D \quad = 1 - \sum_{i=1}^{S} \quad {P_i}^2 \qquad (2)$$

a. where $Pi$ is the relative abundance of avian species $i$.

iv) Evenness (*J*) which measures the relative abundance of different species in a community relative to an ideal condition (Pielou, 1966). A community has high evenness if all species are represented by a similar number of individuals. Conversely, if some species are dominant and others are rare, the evenness is low. Evenness was derived as mosquito Shannon index ($H$ ) divided by the logarithm of mosquito species richness (*S*):

$$J = \frac{H}{ln(S)} \qquad (3)$$

**Section S5: Trait imputation**

The functional identity of the avian community was defined using a matrix of 24 morphological and life-history traits (Table S1) collected from global databases, including AVONET database (Tobias et al., 2022), Amniote database (Myhrvold et al., 2015), and data provided by Tobias and Pigot (2019). We imputed missing values where necessary, using the *missForest* R package (Stekhoven, 2011). To enhance the robustness of our data imputation, we included phylogenetic eigenvectors as predictors (Diniz-Filho et al., 2012; Penone et al., 2014). To determine the optimal number of eigenvectors for improving imputation accuracy, we repeated the procedure using datasets containing the first 5, 10, 15, or 20 eigenvectors and assessed the accuracy of the imputation by measuring the Normalized Root Mean Square Error (NRMSE), where lower values indicate better imputation performance (Figure S2). The imputation procedure was generally successful in filling data gaps (NRMSE $< 0.5$), with the set of 10 eigenvectors proving to be the most accurate. However, we chose to exclude clutches per year, maximum longevity, and incubation from our analysis because their NRMSE values were consistently above 0.5, indicating insufficient imputation accuracy.

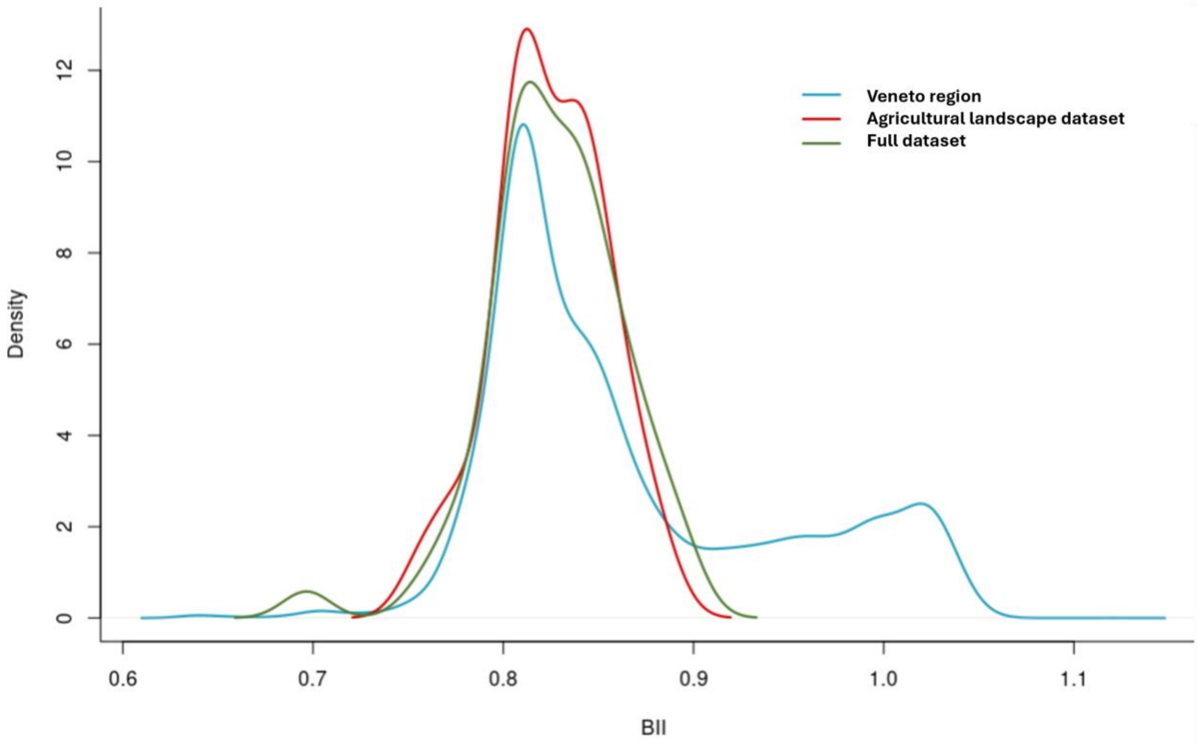

**Figure S1.** Distribution of the Biodiversity Intactness Index (BII; Newbold et al., 2016) values across the Veneto region compared to the sampling coverage of the entomological surveillance network. Density plots show the distribution of BII values for three spatial extents: Veneto (the full geographical extent of the region, blue line); agricultural landscape dataset (the BII values associated with the geographically constrained, empirically sampled FBI trap network, red line); and full dataset (the BII values associated with all trap locations used in the full analysis, green line).

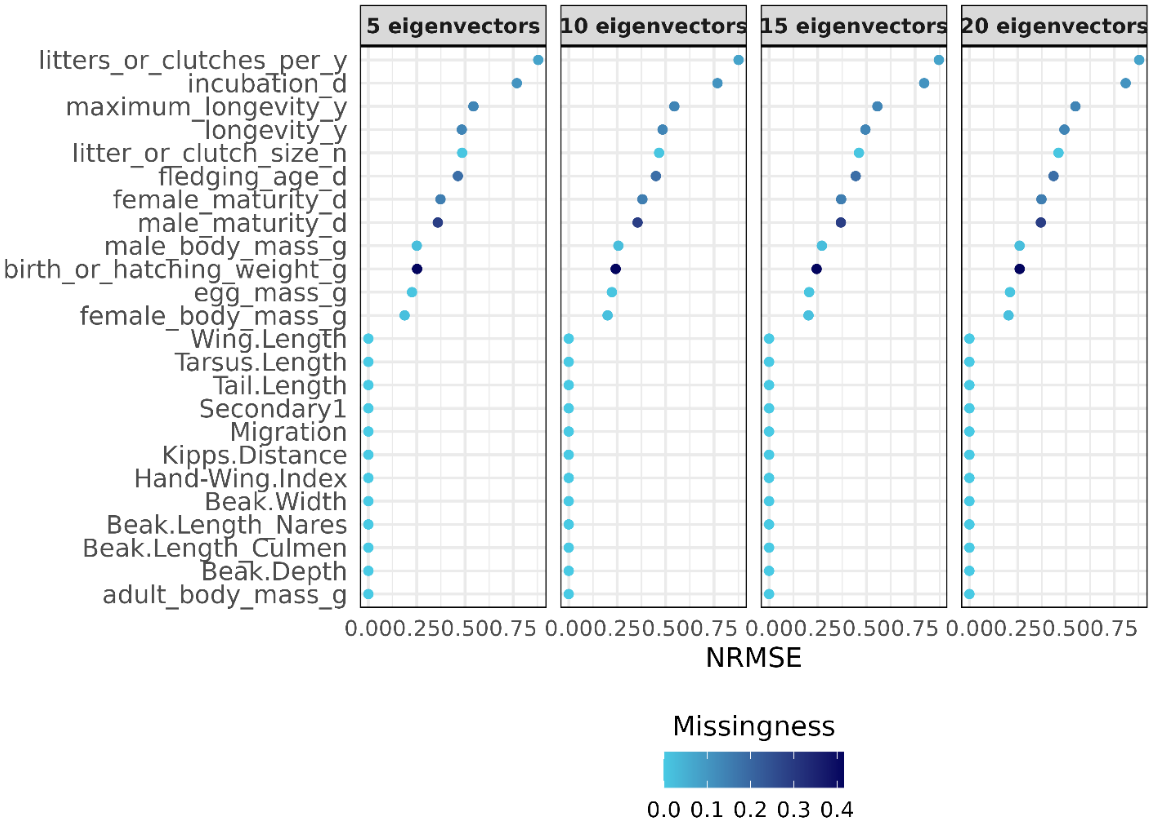

**Figure S2.** Effect of the number of eigenvectors on trait imputation error. Four panels show results for 5, 10, 15, and 20 eigenvectors. For each trait (y-axis), the normalized root mean squared error (NMRSE) is plotted on the x-axis. Point color indicates the proportion of missing data for each trait.

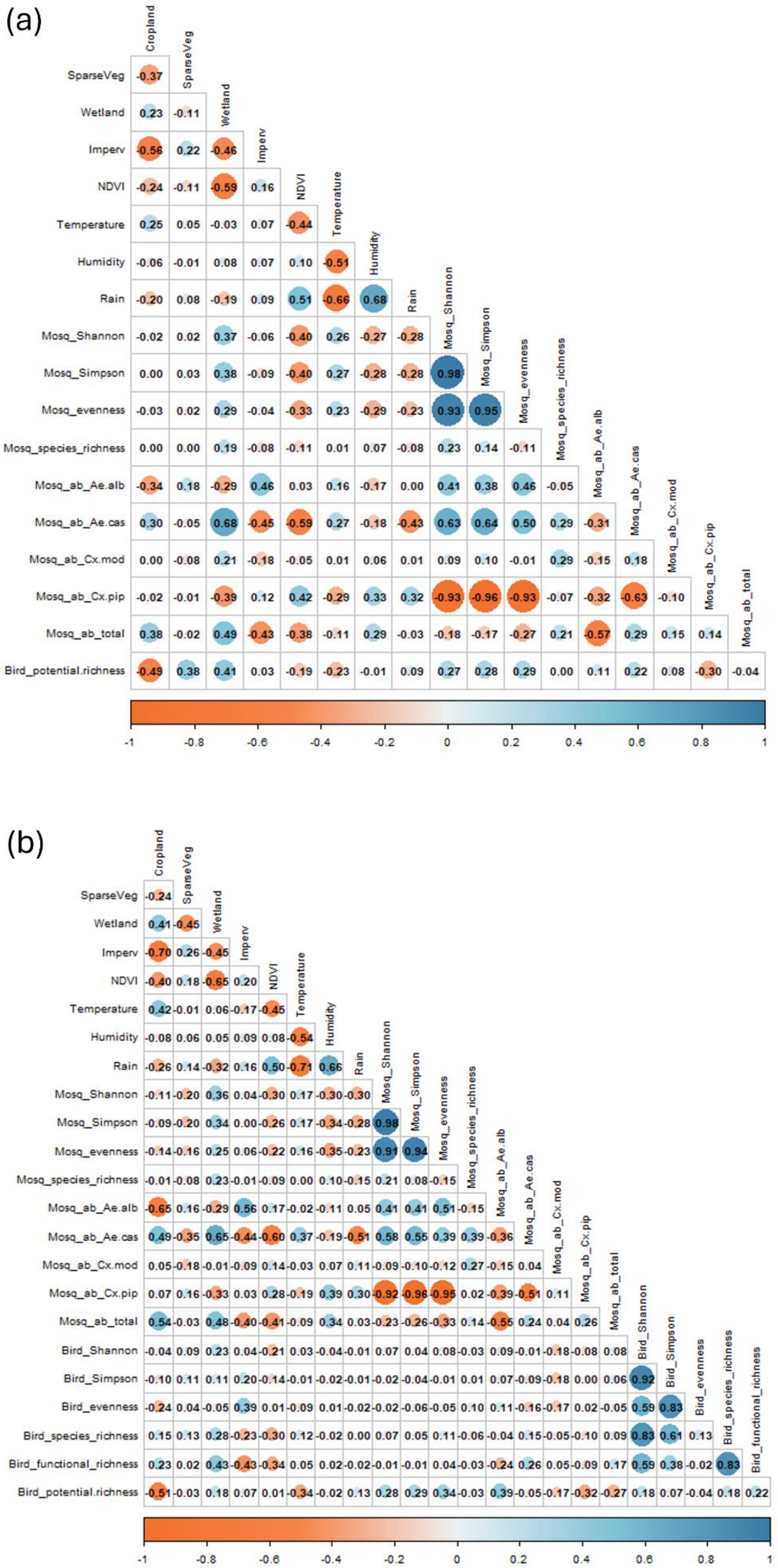

**Figure S3**. Spearman's correlation coefficients among explanatory variables, based on a) the full dataset b) the agricultural landscape dataset.

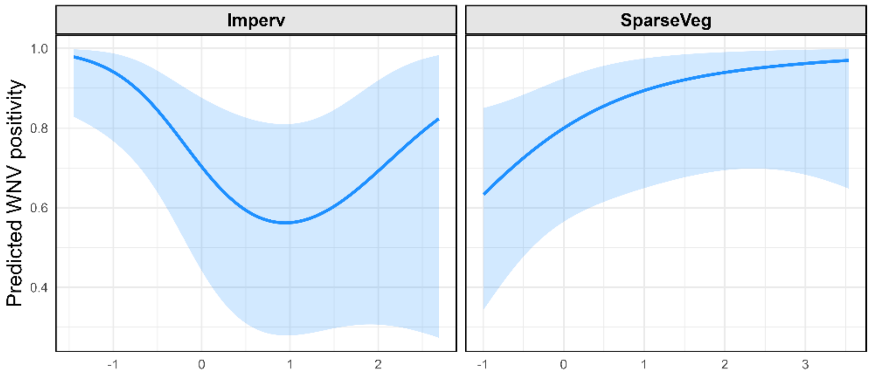

**Figure S4.** Partial dependence plots showing the partial effects of environmental variables on the predicted WNV positivity in mosquito pools. The plots display the predicted probability derived from the minimum adequate environmental model, fitted using the agricultural landscape dataset. Each curve represents the isolated, non-linear effect of the continuous environmental variable (listed on the x-axis, centered and scaled) on WNV positivity, holding all other covariates constant at their median values. The shaded areas indicate the 95% confidence intervals around the predicted effect.

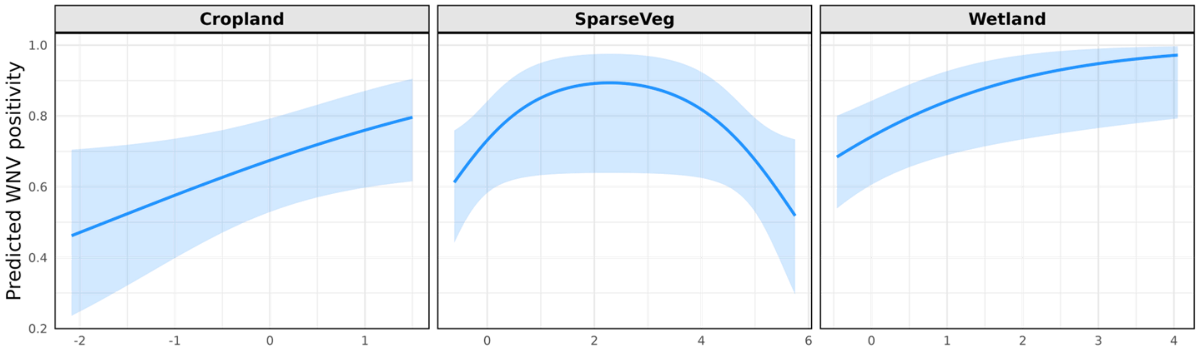

**Figure S5.** Partial dependence plots showing the partial effects of environmental variables on the predicted WNV positivity in mosquito pools. The plots display the predicted probability derived from the minimum adequate environmental model, fitted using the full dataset. Each curve represents the isolated, non-linear effect of the continuous environmental variable (listed on the x-axis, centered and scaled) on WNV positivity, holding all other covariates constant at their median values. The shaded areas indicate the 95% confidence intervals around the predicted effect.

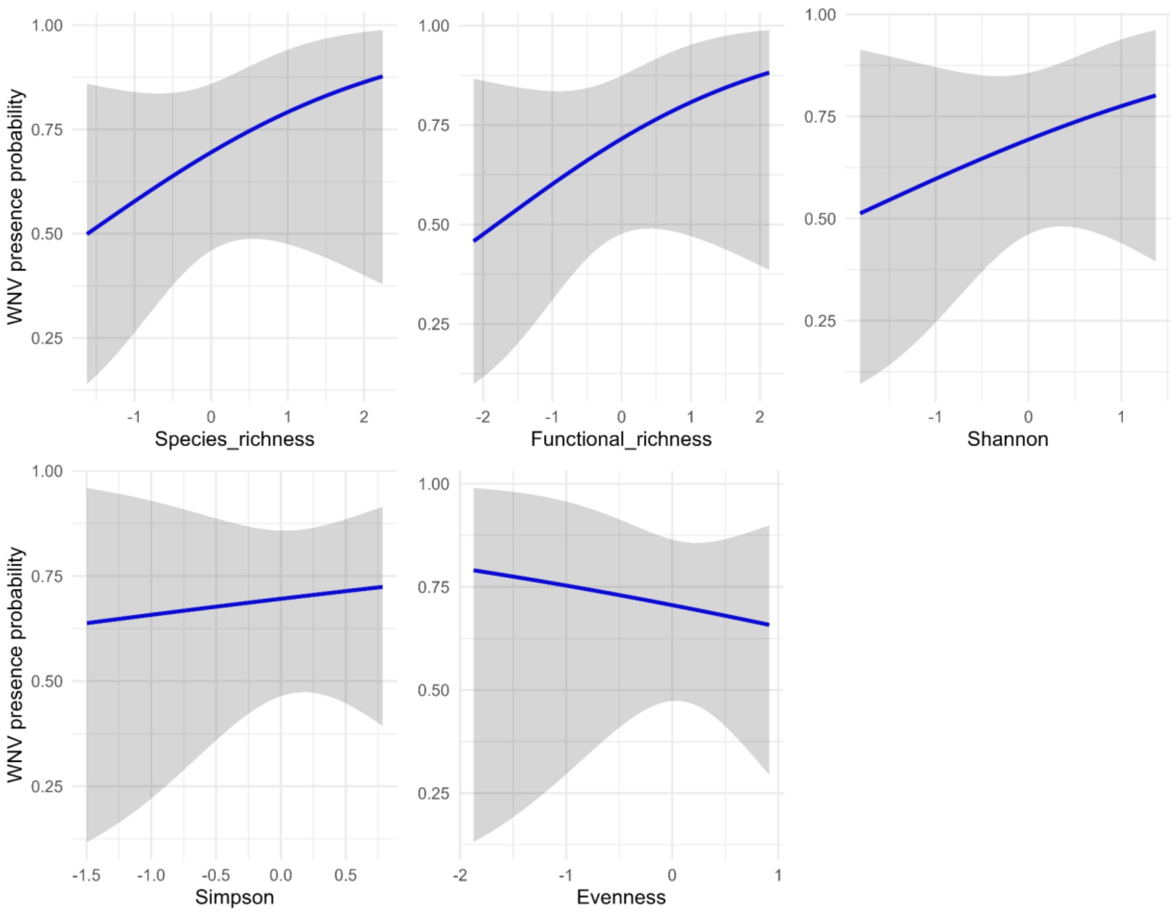

**Figure S6. Partial dependence plots** showing the univariate effects of avian biodiversity metrics on the probability of WNV presence in mosquito pools, fitted on the high-completeness subset (Chao2 sample completeness > 0.5).

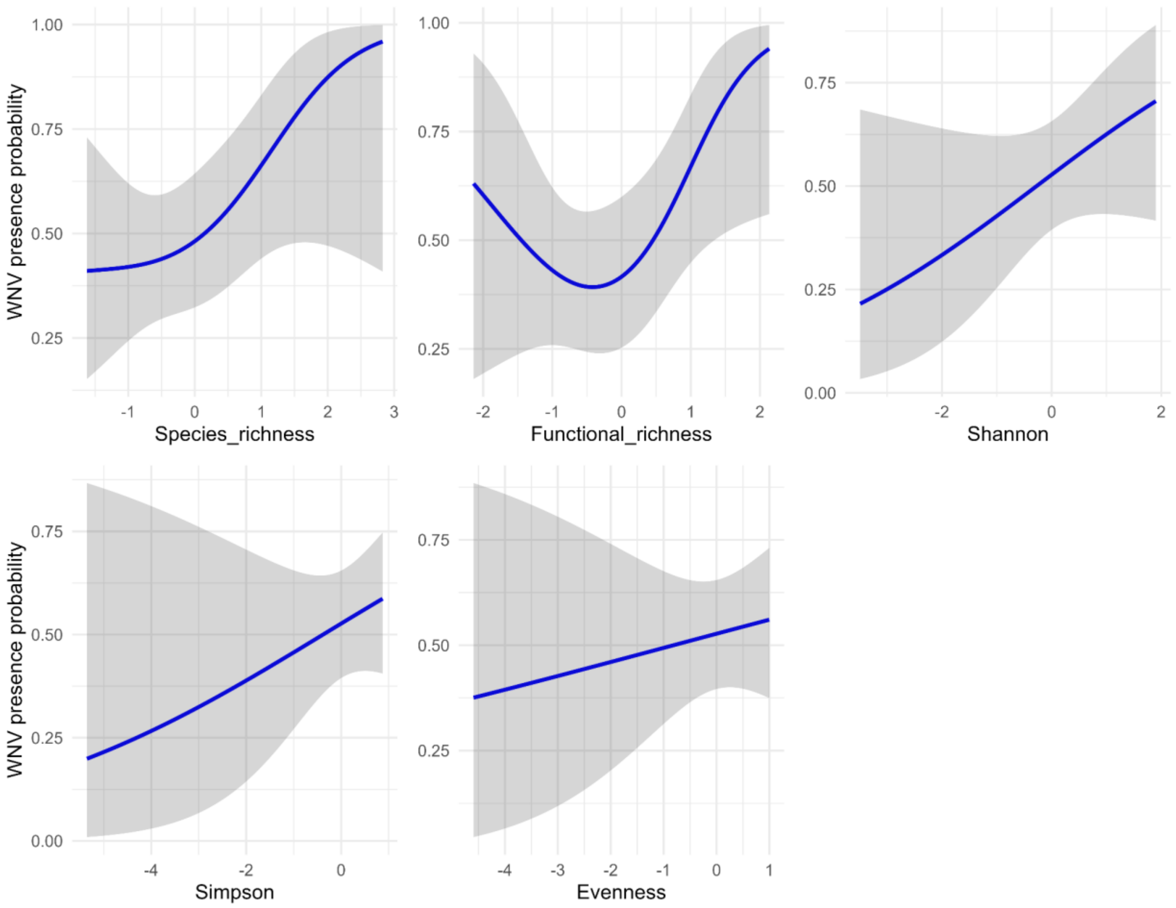

**Figure S7. Partial dependence plots** showing the univariate effects of avian biodiversity metrics on the probability of WNV presence in mosquito pools, fitted on the agricultural landscape dataset (n = 55).

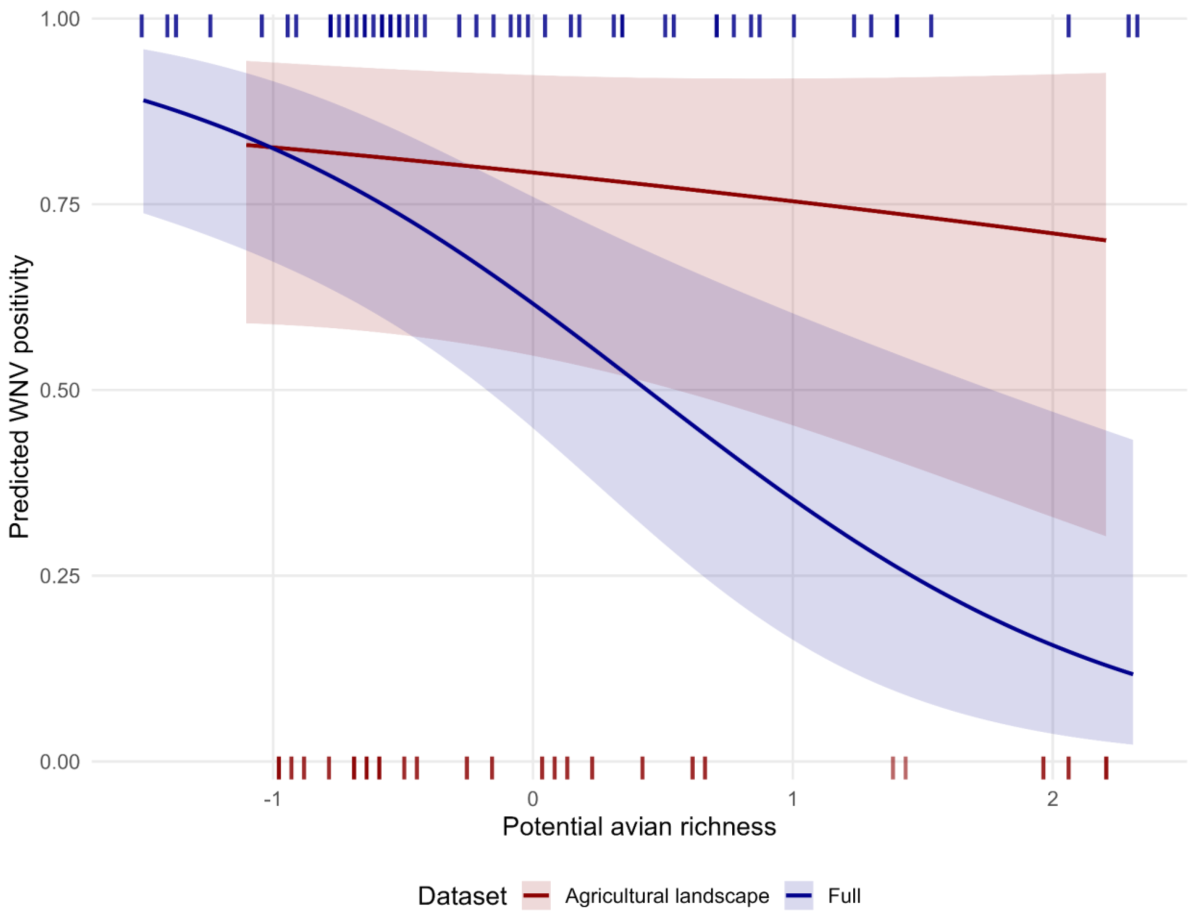

**Figure S8.** Predicted probability of WNV positivity in mosquitoes as a response to potential avian species, using the agricultural landscape dataset and (red) the full dataset (blue). Curves are plotted over the central 95% of observed values for each metric (observation densities are reported at the bottom of each plot). Shaded areas indicate 95% confidence intervals. Values on the x axis have been rescaled for analysis. These results are derived from integrated host-vector models combining avian richness with *Cx. pipiens* relative abundance.

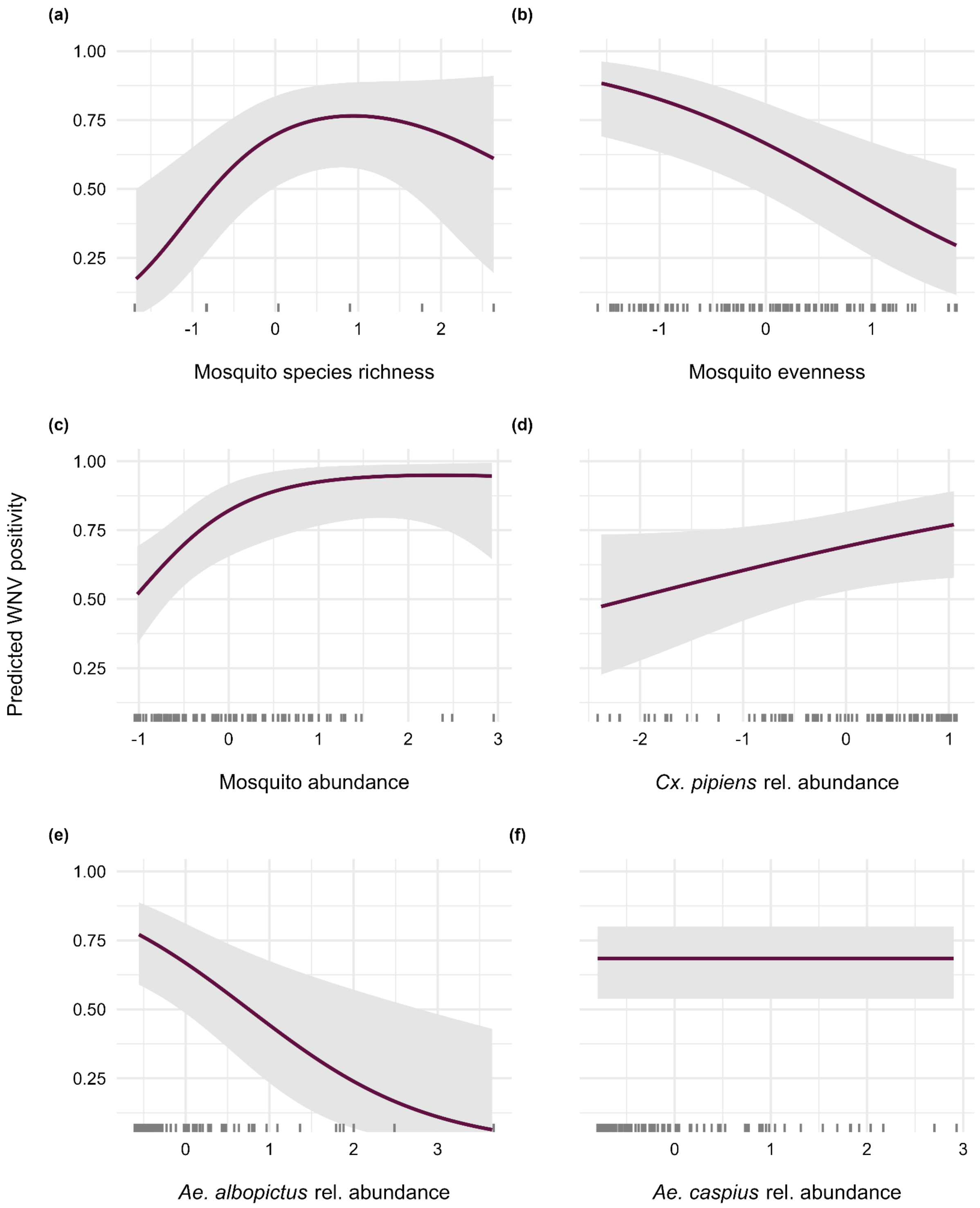

**Figure S9.** Predicted probability of WNV positivity in mosquitoes as a response to different mosquito biodiversity metrics. Curves are plotted over the central 95% of observed values for each metric (observation densities are reported at the bottom of each plot). Shaded grey areas indicate 95% confidence intervals. Values on the x axis have been rescaled for analysis. These results are derived from models fitted on the full dataset.

**Table S1.** Avian traits used to calculate functional richness.

| Trait | Unit | Reference |
|---|---|---|
| Female maturity | day | Myhrvold et al., 2015 |
| Clutch size | num | Myhrvold et al., 2015 |
| Adult body mass | gr | Myhrvold et al., 2015 |
| Birth or hatching weight | gr | Myhrvold et al., 2015 |
| Egg mass | gr | Myhrvold et al., 2015 |
| Fledging age | day | Myhrvold et al., 2015 |
| Maximum longevity | year | Myhrvold et al., 2015 |
| Clutches per year | num | Myhrvold et al., 2015 |
| Incubation | day | Myhrvold et al., 2015 |
| Longevity | year | Myhrvold et al., 2015 |
| Male maturity | day | Myhrvold et al., 2015 |
| Female body mass | gr | Myhrvold et al., 2015 |
| Male body mass | gr | Myhrvold et al., 2015 |
| Beak length (culmen) | mm | Tobias et al., 2022 |
| Beak length (nares) | mm | Tobias et al., 2022 |
| Beak width | mm | Tobias et al., 2022 |
| Beak depth | mm | Tobias et al., 2022 |
| Tarsus length | mm | Tobias et al., 2022 |
| Wing length | mm | Tobias et al., 2022 |
| Kipp’s distance | mm | Tobias et al., 2022 |
| Secondary1 | mm | Tobias et al., 2022 |
| Hand-wing index | index | Tobias et al., 2022 |
| Tail length | mm | Tobias et al., 2022 |
| Migration | 1 = Migratory and partially migratory; 0 = Non migratory | Tobias & Pigot, 2019 |

**Table S2**. Minimum adequate GAMs predicting WNV positivity at the 5 km buffer scale. Models were fitted on the agricultural landscape dataset.

| **Model** | | | | | |
|---|---|---|---|---|---|
| **Environmental1** | | **Estimate** | **Std. Error** | **z value** | **Pr(>\|z\|)** |
| | (Intercept) | 1.87590439 | 0.64537634 | 2.90668293 | 0.00365283 |
| | year2023 | -2.4463215 | 0.91645501 | -2.6693307 | 0.00760026 |
| | | **edf** | **Ref.df** | **Chi.sq** | **p-value** |
| | s(Humidity) | 0.780442176 | 2 | 2.701333338 | 0.059543694 |
| | s(Temperature) | 0.5598006 | 2 | 0.96861663 | 0.177685197 |
| | s(NDVI) | 6.23475E-06 | 2 | 3.39646E-07 | 0.88327455 |
| | s(Cropland) | 2.88374E-05 | 2 | 1.02127E-05 | 0.564855516 |
| | s(Imperv) | 1.476903991 | 2 | 5.143172359 | 0.023818954 |
| | s(SparseVeg) | 1.574674823 | 2 | 7.874875331 | 0.004614634 |
| | s(Wetland) | 0.767555061 | 2 | 1.828314231 | 0.114085571 |
| | ti(Lon.Lat) | 1.36896388 | 16 | 5.570365622 | 0.012669089 |
| | | **$R^2$ (adj.)** = 0.486 | | **Deviance explained** = 46.9% | |
| **Environmental2** | | **Estimate** | **Std. Error** | **z value** | **Pr(>\|z\|)** |
| | (Intercept) | 1.87590493 | 0.64537695 | 2.90668101 | 0.00365285 |
| | year2023 | -2.4463229 | 0.91645683 | -2.669327 | 0.00760034 |
| | | **edf** | **Ref.df** | **Chi.sq** | **p-value** |
| | s(Humidity) | 0.780445548 | 2 | 2.70135253 | 0.059543196 |
| | s(Temperature) | 0.559802291 | 2 | 0.968617297 | 0.177685431 |
| | s(Cropland) | 1.24859E-05 | 2 | 3.13322E-06 | 0.667136762 |
| | s(Imperv) | 1.476905399 | 2 | 5.143259514 | 0.023818283 |
| | s(SparseVeg) | 1.574674586 | 2 | 7.87485949 | 0.004614663 |
| | s(Wetland) | 0.767558287 | 2 | 1.828314531 | 0.114086055 |
| | ti(Lon.Lat) | 1.368969282 | 16 | 5.570368104 | 0.012669105 |
| | | **$R^2$ (adj.)** = 0.486 | | **Deviance explained** = 46.9% | |
| **Environmental3** | | **Estimate** | **Std. Error** | **z value** | **Pr(>\|z\|)** |
| | (Intercept) | -1.140533509 | 0.378133767 | -3.016217037 | 0.0025595 |
| | year2023 | 2.204918634 | 0.548376061 | 4.020814893 | 5.79972E-05 |

| | | edf | Ref.df | Chi.sq | p-value |
|---|---|---|---|---|---|
| | s(Humidity) | 0.78044525 | 2 | 2.70135267 | 0.05954321 |
| | s(Temperature) | 0.55980343 | 2 | 0.96861551 | 0.17768622 |
| | s(Imperv) | 1.47690675 | 2 | 5.14328353 | 0.0238182 |
| | s(SparseVeg) | 1.57467492 | 2 | 7.87487732 | 0.00461463 |
| | s(Wetland) | 0.76755569 | 2 | 1.82831356 | 0.11408594 |
| | ti(Lon.Lat) | 1.36896715 | 16 | 5.5703712 | 0.01266908 |
| | | **$R^2$ (adj.)** = 0.486 | | **Deviance explained** = 46.9% | |
| **Environmental4** | | **Estimate** | **Std. Error** | **z value** | **Pr(>\|z\|)** |
| | (Intercept) | 1.64696497 | 0.60081146 | 2.7412343 | 0.00612088 |
| | year2023 | -2.0661582 | 0.80612688 | -2.5630683 | 0.01037516 |
| | | **edf** | **Ref.df** | **Chi.sq** | **p-value** |
| | s(Humidity) | 0.76058468 | 2 | 2.427847869 | 0.07076242 |
| | s(Imperv) | 1.500435977 | 2 | 4.77874199 | 0.034449 |
| | s(SparseVeg) | 1.512416568 | 2 | 7.511905243 | 0.004976571 |
| | s(Wetland) | 0.793351135 | 2 | 2.079496176 | 0.096941591 |
| | ti(Lon.Lat) | 1.313897187 | 16 | 5.007478347 | 0.017181404 |
| | | **$R^2$ (adj.)** = 0.461 | | **Deviance explained** = 44.5% | |
| **Environmental5** | | **Estimate** | **Std. Error** | **z value** | **Pr(>\|z\|)** |
| | (Intercept) | 1.572029 | 0.58288931 | 2.69695971 | 0.00699757 |
| | year2023 | -2.0296484 | 0.77366624 | -2.6234161 | 0.00870529 |
| | | **edf** | **Ref.df** | **Chi.sq** | **p-value** |
| | s(Humidity) | 0.726453726 | 2 | 2.101350782 | 0.085508542 |
| | s(Imperv) | 1.684776743 | 2 | 7.935337522 | 0.007150861 |
| | s(SparseVeg) | 1.165267954 | 2 | 5.832303038 | 0.00843023 |
| | ti(Lon.Lat) | 1.142324549 | 16 | 4.7248401 | 0.017523907 |
| | | **R2 (adj.)** = 0.406 | | **Deviance explained** = 39.5% | |
| **Environmental6** | | **Estimate** | **Std. Error** | **z value** | **Pr(>\|z\|)** |
| | (Intercept) | 1.6105873 | 0.54492901 | 2.95559103 | 0.00312071 |
| | year2023 | -2.3635702 | 0.7015732 | -3.3689574 | 0.00075453 |

| | | | | | |
|---|---|---|---|---|---|
| | | **edf** | **Ref.df** | **Chi.sq** | **p-value** |
| | s(Imperv) | 1.649398152 | 2 | 7.137067241 | 0.011477116 |
| | s(SparseVeg) | 1.015300905 | 2 | 4.222793566 | 0.024635205 |
| | ti(Lon.Lat) | 0.787446064 | 16 | 3.343545129 | 0.034260207 |
| | | **R2 (adj.)** = 0.337 | | **Deviance explained** = 33.7% | |

**Table S3**. Minimum adequate GAMs predicting WNV positivity at the 10 km buffer scale. Models were fitted on the agricultural landscape dataset.

| **Model** | | | | | |
|---|---|---|---|---|---|
| **Environmental1** | | **Estimate** | **Std. Error** | **z value** | **Pr(>\|z\|)** |
| | (Intercept) | -1.140533987 | 0.378135469 | -3.016204719 | 0.002559604 |
| | year2023 | 2.204920329 | 0.548380764 | 4.020783507 | 5.80049E-05 |
| | | **edf** | **Ref.df** | **Chi.sq** | **p-value** |
| | s(Humidity) | 1.64103E-05 | 2 | 1.08969E-05 | 0.435473897 |
| | s(Temperature) | 2.16268E-05 | 2 | 1.34201E-05 | 0.436564809 |
| | s(Rain) | 1.42255E-05 | 2 | 8.38899E-06 | 0.453704297 |
| | s(NDVI) | 0.517325203 | 2 | 0.928866906 | 0.160495985 |
| | s(Cropland) | 0.790897047 | 2 | 3.37888012 | 0.029281132 |
| | s(Imperv) | 0.9206715 | 2 | 10.03594116 | 0.000488145 |
| | s(SparseVeg) | 3.41295E-06 | 2 | 6.29325E-07 | 0.747180983 |
| | s(Wetland) | 0.968828438 | 2 | 2.393878324 | 0.085097205 |
| | ti(Lon.Lat) | 0.98243494 | 16 | 2.145978798 | 0.096402651 |
| | | **$R^2$ (adj.)** = 0.340 | | **Deviance explained** = 31.3% | |
| **Environmental2** | | **Estimate** | **Std. Error** | **z value** | **Pr(>\|z\|)** |
| | (Intercept) | -1.110876112 | 0.374083958 | -2.969590353 | 0.002981971 |
| | year2023 | 2.149247291 | 0.542199782 | 3.963939794 | 7.37229E-05 |
| | | **edf** | **Ref.df** | **Chi.sq** | **p-value** |
| | s(Humidity) | 3.51656E-05 | 2 | 2.38083E-05 | 0.43029897 |
| | s(Temperature) | 2.07137E-05 | 2 | 9.9553E-06 | 0.528952884 |

| | | | | | |
|---|---|---|---|---|---|
| | s(Rain) | 2.35686E-05 | 2 | 1.10811E-05 | 0.564671963 |
| | s(NDVI) | 0.384410897 | 2 | 0.559651815 | 0.209385866 |
| | s(Cropland) | 0.764478322 | 2 | 2.928950455 | 0.039917147 |
| | s(Imperv) | 0.917117815 | 2 | 9.585146228 | 0.00074659 |
| | s(Wetland) | 0.692790518 | 2 | 1.97510052 | 0.086353318 |
| | ti(Lon.Lat) | 1.022790657 | 16 | 2.342049751 | 0.087982445 |
| | | **$R^2$ (adj.)** = 0.332 | | **Deviance explained** = 30.3% | |
| **Environmental3** | | **Estimate** | **Std. Error** | **z value** | **Pr(>\|z\|)** |
| | (Intercept) | -1.140533509 | 0.378133767 | -3.016217037 | 0.0025595 |
| | year2023 | 2.204918634 | 0.548376061 | 4.020814893 | 5.79972E-05 |
| | | **edf** | **Ref.df** | **Chi.sq** | **p-value** |
| | s(Humidity) | 3.51608E-05 | 2 | 2.17642E-05 | 0.477594176 |
| | s(Temperature) | 6.78494E-06 | 2 | 1.33116E-06 | 0.776555032 |
| | s(NDVI) | 0.517328553 | 2 | 0.928862101 | 0.160497813 |
| | s(Cropland) | 0.790902302 | 2 | 3.378875856 | 0.029281287 |
| | s(Imperv) | 0.920676478 | 2 | 10.03594135 | 0.000488145 |
| | s(Wetland) | 0.96882816 | 2 | 2.393883941 | 0.085096454 |
| | ti(Lon.Lat) | 0.98243018 | 16 | 2.14597022 | 0.096403136 |
| | | **$R^2$ (adj.)** = 0.340 | | **Deviance explained** = 31.3% | |
| **Environmental4** | | **Estimate** | **Std. Error** | **z value** | **Pr(>\|z\|)** |
| | (Intercept) | -1.140533423 | 0.378133597 | -3.016218164 | 0.00255949 |
| | year2023 | 2.204918568 | 0.548375496 | 4.020818919 | 5.79962E-05 |

| | | | | | |
|---|---|---|---|---|---|
| | | **edf** | **Ref.df** | **Chi.sq** | **p-value** |
| | s(Humidity) | 3.4025E-05 | 2 | 2.05034E-05 | 0.493397791 |
| | s(NDVI) | 0.517331161 | 2 | 0.928863157 | 0.160498061 |
| | s(Cropland) | 0.790903059 | 2 | 3.378884091 | 0.029281337 |
| | s(Imperv) | 0.920675371 | 2 | 10.03592932 | 0.000488103 |
| | s(Wetland) | 0.968827626 | 2 | 2.393879297 | 0.085096282 |
| | ti(Lon.Lat) | 0.982440805 | 16 | 2.145974068 | 0.096403902 |
| | | **$R^2$ (adj.)** = 0.340 | | **Deviance explained** = 31.3% | |
| **Environmental5** | | **Estimate** | **Std. Error** | **z value** | **Pr(>\|z\|)** |
| | (Intercept) | -1.140538321 | 0.3781299 | -3.016260602 | 0.002559132 |
| | year2023 | 2.204929674 | 0.548363832 | 4.020924697 | 5.79701E-05 |
| | | **edf** | **Ref.df** | **Chi.sq** | **p-value** |
| | s(NDVI) | 0.517327847 | 2 | 0.928867957 | 0.1604968 |
| | s(Cropland) | 0.790899168 | 2 | 3.378918678 | 0.029281075 |
| | s(Imperv) | 0.920670263 | 2 | 10.03592125 | 0.000488106 |
| | s(Wetland) | 0.968828864 | 2 | 2.393885684 | 0.085096721 |
| | ti(Lon.Lat) | 0.982450087 | 16 | 2.145974882 | 0.096405096 |
| | | **R2 (adj.)** = 0.340 | | **Deviance explained** = 31.3% | |
| **Environmental6** | | **Estimate** | **Std. Error** | **z value** | **Pr(>\|z\|)** |
| | (Intercept) | -1.068011882 | 0.36697838 | -2.910285568 | 0.003610987 |
| | year2023 | 2.085364593 | 0.536274974 | 3.888610683 | 0.00010082 |

| | | | | | |
|---|---|---|---|---|---|
| | | edf | Ref.df | Chi.sq | p-value |
| | s(Cropland) | 0.701180971 | 2 | 2.155086096 | 0.076727539 |
| | s(Imperv) | 0.912939938 | 2 | 9.190621321 | 0.000993336 |
| | s(Wetland) | 0.695512619 | 2 | 2.011113974 | 0.083806813 |
| | ti(Lon.Lat) | 1.009308798 | 16 | 2.27117867 | 0.092264203 |
| | | **R2 (adj.)** = 0.324 | | **Deviance explained** = 29.2% | |
| **Environmental7** | | **Estimate** | **Std. Error** | **z value** | **Pr(>\|z\|)** |
| | (Intercept) | -1.0367512 | 0.35706152 | -2.9035646 | 0.00368941 |
| | year2023 | 2.04834104 | 0.52719119 | 3.88538558 | 0.00010217 |
| | | **edf** | **Ref.df** | **Chi.sq** | **p-value** |
| | s(Cropland) | 0.614799079 | 2 | 1.498310251 | 0.11633519 |
| | s(Imperv) | 0.914028573 | 2 | 9.440504547 | 0.000808129 |
| | ti(Lon.Lat) | 0.742659753 | 16 | 1.286796612 | 0.164796977 |
| | | **R2 (adj.)** = 0.298 | | **Deviance explained** = 26.2% | |
| **Environmental8** | | **Estimate** | **Std. Error** | **z value** | **Pr(>\|z\|)** |
| | (Intercept) | -1.041553 | 0.35786051 | -2.9104999 | 0.00360851 |
| | year2023 | 2.03105739 | 0.52576551 | 3.86304801 | 0.00011198 |
| | | **edf** | **Ref.df** | **Chi.sq** | **p-value** |
| | s(Imperv) | 1.390062743 | 2 | 12.01988741 | 0.000269435 |
| | ti(Lon.Lat) | 0.869231319 | 16 | 1.647060723 | 0.133720407 |

| | | | | |
|---|---|---|---|---|
| | | **R2 (adj.)** = 0.294 | | **Deviance explained** = 25.9% |

**Table S4.** Minimum adequate GAMs predicting WNV positivity at the 5 km buffer scale. Models were fitted on the full dataset.

| Model | | | | | |
|---|---|---|---|---|---|
| **Environmental1** | | **Estimate** | **Std. Error** | **z value** | **Pr(>\|z\|)** |
| | (Intercept) | 1.09609719 | 0.31970972 | 3.42841367 | 0.00060712 |
| | year2023 | -1.9677515 | 0.45259971 | -4.347664 | 1.376E-05 |
| | | **edf** | **Ref.df** | **Chi.sq** | **p-value** |
| | s(Humidity) | 4.76363E-06 | 2 | 1.2226E-06 | 0.772207314 |
| | s(Temperature) | 8.72261E-06 | 2 | 5.68117E-06 | 0.474392357 |
| | s(NDVI) | 3.84129E-06 | 2 | 9.3016E-09 | 0.996934406 |
| | s(Cropland) | 0.813917949 | 2 | 4.078347791 | 0.023984857 |
| | s(Imperv) | 0.636968236 | 2 | 1.649114843 | 0.106048282 |
| | s(SparseVeg) | 0.805268097 | 2 | 3.787102302 | 0.028548321 |
| | s(Wetland) | 0.848087977 | 2 | 4.587135701 | 0.018938886 |
| | ti(Lon.Lat) | 3.28417E-05 | 16 | 1.17268E-05 | 0.854531985 |
| | | **$R^2$ (adj.)** = 0.284 | | **Deviance explained** = 24.7% | |
| **Environmental2** | | **Estimate** | **Std. Error** | **z value** | **Pr(>\|z\|)** |
| | (Intercept) | 1.09609336 | 0.31971099 | 3.42838815 | 0.00060718 |
| | year2023 | -1.9677446 | 0.45260296 | -4.3476177 | 1.3762E-05 |
| | | **edf** | **Ref.df** | **Chi.sq** | **p-value** |
| | s(Humidity) | 6.02623E-06 | 2 | 1.56836E-06 | 0.769461144 |
| | s(Temperature) | 3.64793E-05 | 2 | 2.66246E-05 | 0.404455085 |
| | s(Cropland) | 0.813916863 | 2 | 4.0782853 | 0.023985437 |

| | | | | | |
|---|---|---|---|---|---|
| | s(Imperv) | 0.636972505 | 2 | 1.649123353 | 0.106047593 |
| | s(SparseVeg) | 0.805268353 | 2 | 3.787074853 | 0.028548721 |
| | s(Wetland) | 0.848088478 | 2 | 4.587134187 | 0.018938989 |
| | ti(Lon.Lat) | 3.05315E-05 | 16 | 1.03694E-05 | 0.900537712 |
| | | **$R^2$ (adj.)** = 0.284 | | **Deviance explained** = 24.7% | |
| **Environmental3** | | **Estimate** | **Std. Error** | **z value** | **Pr(>\|z\|)** |
| | (Intercept) | 1.09609445 | 0.319709765 | 3.428404666 | 0.00060714 |
| | year2023 | -1.96774668 | 0.452599569 | -4.347654777 | 1.37601E-05 |
| | | **edf** | **Ref.df** | **Chi.sq** | **p-value** |
| | s(Temperature) | 3.00224E-05 | 2 | 2.1046E-05 | 0.429778549 |
| | s(Cropland) | 0.813915691 | 2 | 4.078292525 | 0.023985305 |
| | s(Imperv) | 0.636974141 | 2 | 1.649122013 | 0.106047993 |
| | s(SparseVeg) | 0.805268887 | 2 | 3.787084013 | 0.028548614 |
| | s(Wetland) | 0.848088083 | 2 | 4.587132407 | 0.018938988 |
| | ti(Lon.Lat) | 2.95785E-05 | 16 | 1.01468E-05 | 0.895505923 |
| | | **$R^2$ (adj.)** = 0.284 | | **Deviance explained** = 24.7% | |
| **Environmental4** | | **Estimate** | **Std. Error** | **z value** | **Pr(>\|z\|)** |
| | (Intercept) | 1.096098019 | 0.319708823 | 3.428425933 | 0.000607092 |
| | year2023 | -1.967753248 | 0.452596859 | -4.347695323 | 1.37576E-05 |
| | | **edf** | **Ref.df** | **Chi.sq** | **p-value** |
| | s(Cropland) | 0.813914619 | 2 | 4.078333704 | 0.023984844 |

| | | | | | |
|---|---|---|---|---|---|
| | s(Imperv) | 0.636974872 | 2 | 1.649113767 | 0.106049324 |
| | s(SparseVeg) | 0.805269558 | 2 | 3.787116751 | 0.028548191 |
| | s(Wetland) | 0.848088195 | 2 | 4.58713017 | 0.018939029 |
| | ti(Lon.Lat) | 3.68642E-05 | 16 | 9.71588E-06 | 0.947284168 |
| | | **$R^2$ (adj.)** = 0.284 | | **Deviance explained** = 24.7% | |
| **Environmental5** | | **Estimate** | **Std. Error** | **z value** | **Pr(>\|z\|)** |
| | (Intercept) | 1.08617339 | 0.31829832 | 3.41243837 | 0.00064384 |
| | year2023 | -1.948511 | 0.44969913 | -4.3329215 | 1.4714E-05 |
| | | **edf** | **Ref.df** | **Chi.sq** | **p-value** |
| | s(Cropland) | 0.812487473 | 2 | 4.100126009 | 0.023495298 |
| | s(SparseVeg) | 0.793246274 | 2 | 3.545864292 | 0.032958713 |
| | s(Wetland) | 0.879272463 | 2 | 6.066815871 | 0.00853295 |
| | ti(Lon.Lat) | 6.86494E-05 | 16 | 1.75087E-05 | 0.956606013 |
| | | **$R^2$ (adj.)** = 0.269 | | **Deviance explained** = 23.3% | |

**Table S5.** Summary results of GAMs testing the effect of different avian biodiversity predictors on the probability of detecting WNV in mosquito pools. Models were fitted on the agricultural landscape dataset.

| **Model** | | | | | |
|---|---|---|---|---|---|
| **Avian species richness** | | **Estimate** | **Std. Error** | **z value** | **Pr(>\|z\|)** |
| | (Intercept) | 2.073757731 | 0.68894618 | 3.010043152 | 0.002612106 |
| | year2023 | -3.487748678 | 0.973467842 | -3.582808315 | 0.00033992 |
| | points | -3.251088658 | 1.058517036 | -3.071361676 | 0.002130849 |
| | | **edf** | **Ref.df** | **Chi.sq** | **p-value** |
| | s(Species_richness) | 1.737601801 | 2 | 10.84251824 | 0.002019142 |
| | s(Imperv) | 3.72002E-06 | 2 | 1.99085E-07 | 0.93753582 |
| | s(SparseVeg) | 0.797701711 | 2 | 3.309066121 | 0.038473933 |
| | ti(Lon.Lat) | 0.443835377 | 15 | 0.780323331 | 0.179925202 |
| | | **$R^2$ (adj.)** = 0.558 | | **Deviance explained** = 54.6% | |
| **Avian functional richness** | | **Estimate** | **Std. Error** | **z value** | **Pr(>\|z\|)** |
| | (Intercept) | 1.557368078 | 0.54165646 | 2.875195246 | 0.004037777 |
| | year2023 | -2.504080911 | 0.739717914 | -3.385183543 | 0.000711307 |
| | points | -0.965895647 | 0.467972341 | -2.064001573 | 0.039017567 |
| | | **edf** | **Ref.df** | **Chi.sq** | **p-value** |
| | s(Functional_richness) | 1.728110052 | 2 | 8.525525773 | 0.006549615 |
| | s(Imperv) | 1.75756E-05 | 2 | 1.14925E-05 | 0.407783967 |
| | s(SparseVeg) | 1.287887517 | 2 | 4.268463788 | 0.034827817 |
| | ti(Lon.Lat) | 0.647289554 | 16 | 1.705569824 | 0.099707274 |
| | | **$R^2$ (adj.)** = 0.393 | | **Deviance explained** = 38.6% | |
| | | **Estimate** | **Std. Error** | **z value** | **Pr(>\|z\|)** |

| | | | | | |
|---|---|---|---|---|---|
| **Avian Shannon index** | (Intercept) | 2.405994109 | 0.731448362 | 3.289356071 | 0.001004169 |
| | year2023 | -3.324089061 | 0.926914658 | -3.586186747 | 0.000335549 |
| | points | -1.743048655 | 0.682832096 | -2.552675342 | 0.010689909 |
| | | **edf** | **Ref.df** | **Chi.sq** | **p-value** |
| | s(Shannon) | 1.678992721 | 2 | 7.681118383 | 0.009617978 |
| | s(Imperv) | 1.521086843 | 2 | 5.54700743 | 0.020249306 |
| | s(SparseVeg) | 1.516272915 | 2 | 5.670527358 | 0.016479225 |
| | ti(Lon.Lat) | 1.483498196 | 16 | 5.041102366 | 0.016880879 |
| | | **$R^2$ (adj.)** = 0.493 | | **Deviance explained** = 50.8% | |
| ***A*vian Simpson index** | | **Estimate** | **Std. Error** | **z value** | **Pr(>\|z\|)** |
| | (Intercept) | 1.776724025 | 0.580865177 | 3.058754587 | 0.002222591 |
| | year2023 | -2.508000429 | 0.743348234 | -3.373923974 | 0.000741048 |
| | points | -0.462290561 | 0.392685311 | -1.177254531 | 0.239093929 |
| | | **edf** | **Ref.df** | **Chi.sq** | **p-value** |
| | s(Simpson) | 0.7345475 | 2 | 1.876076453 | 0.107250188 |
| | s(Imperv) | 1.733193415 | 2 | 8.617956346 | 0.00557075 |
| | s(SparseVeg) | 1.33742873 | 2 | 5.019635329 | 0.023196505 |
| | ti(Lon.Lat) | 0.849034884 | 16 | 4.574574065 | 0.016502537 |
| | | **$R^2$ (adj.)** = 0.387 | | **Deviance explained** = 40.2% | |
| ***A*vian evenness** | | **Estimate** | **Std. Error** | **z value** | **Pr(>\|z\|)** |
| | (Intercept) | 1.837644362 | 0.592424292 | 3.101905824 | 0.001922791 |
| | year2023 | -2.515484014 | 0.745708727 | -3.373279572 | 0.000742785 |
| | points | -0.33440722 | 0.362257901 | -0.923119191 | 0.355945095 |
| | | **edf** | **Ref.df** | **Chi.sq** | **p-value** |
| | s(Evenness) | 0.718224712 | 2 | 1.831756507 | 0.104230246 |

| | | | | | |
|---|---|---|---|---|---|
| | s(Imperv) | 1.761666132 | 2 | 9.395166864 | 0.003535584 |
| | s(SparseVeg) | 1.324761688 | 2 | 5.786079556 | 0.01295154 |
| | ti(Lon.Lat) | 0.928354757 | 16 | 5.026527844 | 0.012852446 |
| | | **$R^2$ (adj.)** = 0.389 | | **Deviance explained** = 40.5% | |
| **Potential avian species richness** | | **Estimate** | **Std. Error** | **z value** | **Pr(>\|z\|)** |
| | (Intercept) | 1.629387943 | 0.549784742 | 2.963683452 | 0.003039808 |
| | year2023 | -2.399188275 | 0.709289406 | -3.382523769 | 0.000718231 |
| | | **edf** | **Ref.df** | **Chi.sq** | **p-value** |
| | s(pot.richness) | 0.579228148 | 2 | 1.24139987 | 0.137487177 |
| | s(SparseVeg) | 0.825442291 | 2 | 3.885262197 | 0.027046567 |
| | s(Imperv) | 1.639977723 | 2 | 7.60022853 | 0.007315667 |
| | ti(Lon.Lat) | 0.780505702 | 16 | 3.135048254 | 0.039881422 |
| | | **$R^2$ (adj.)** = 0.358 | | **Deviance explained** = 35.5% | |

**Table S6.** Validation of GAMs testing the effect of different avian biodiversity predictors on the probability of detecting WNV in mosquito pools. Models were fitted on the agricultural landscape dataset.

| **Biodiversity variable** | **TPR** | **TNR** | **TSS** |
|---|---|---|---|
| Avian species richness | 0.7931034 | 0.6923077 | 0.4854111 |
| Avian functional richness | 0.7586207 | 0.6538462 | 0.4124668 |
| Avian Shannon index | 0.6896552 | 0.6538462 | 0.3435013 |
| Avian Simpson index | 0.6896552 | 0.5769231 | 0.2665782 |
| Avian evenness | 0.6896552 | 0.6153846 | 0.3050398 |
| Avian potential species richness | 0.7241379 | 0.6153846 | 0.3395225 |

**Table S7.** Summary results of GAMs testing the effect of potential avian species richness alone or combined with *Cx. pipiens* relative abundance on the probability of detecting WNV in mosquito pools. Models were fitted on the full dataset.

| **Model** | | | | | |
|---|---|---|---|---|---|
| **Potential avian species richness** | | **Estimate** | **Std. Error** | **z value** | **Pr(>\|z\|)** |
| | (Intercept) | 1.2205719 | 0.343000608 | 3.558512348 | 0.000372961 |
| | year2023 | -2.129958321 | 0.480435987 | -4.433386298 | 9.27644E-06 |
| | | **edf** | **Ref.df** | **Chi.sq** | **p-value** |
| | s(pot.richness) | 0.92811375 | 2 | 11.47166626 | 0.000322473 |
| | s(SparseVeg) | 0.877516177 | 2 | 6.408768747 | 0.006530507 |
| | s(Cropland) | 4.35132E-05 | 2 | 4.04803E-05 | 0.33430813 |
| | s(Wetland) | 0.933531902 | 2 | 11.66593425 | 0.000336963 |
| | ti(Lon.Lat) | 7.7126E-05 | 16 | 3.44315E-05 | 0.823684706 |
| | | **$R^2$ (adj.)** = 0.341 | | **Deviance explained** = 29.7% | |
| ***Cx. pipiens* relative abundance + potential avian species richness** | | **Estimate** | **Std. Error** | **z value** | **Pr(>\|z\|)** |
| | (Intercept) | 1.248167755 | 0.347685124 | 3.589937186 | 0.000330758 |
| | year2023 | -2.179685891 | 0.490953892 | -4.439695718 | 9.00862E-06 |
| | | **edf** | **Ref.df** | **Chi.sq** | **p-value** |
| | s(Prop_cxpip) | 0.218128535 | 2 | 0.266137066 | 0.268027436 |
| | s(pot.richness) | 0.925066419 | 2 | 10.93009576 | 0.0004077 |
| | s(SparseVeg) | 0.876071926 | 2 | 6.318020325 | 0.006855371 |
| | s(Cropland) | 0.000218187 | 2 | 0.000234111 | 0.296671767 |
| | s(Wetland) | 0.933608445 | 2 | 11.6726515 | 0.000331007 |
| | ti(Lon.Lat) | 4.687E-05 | 16 | 2.06657E-05 | 0.827947407 |
| | | **$R^2$ (adj.)** = 0.342 | | **Deviance explained** = 29.9% | |

**Table S8.** Validation of GAMs testing the effect of potential avian species richness alone or combined with *Cx. pipiens* relative abundance on the probability of detecting WNV in mosquito pools. Models were fitted on the full dataset.

| **Biodiversity variable** | **TPR** | **TNR** | **TSS** |
|---|---|---|---|
| Potential avian species richness | 0.779661 | 0.7636364 | 0.5432974 |
| *Cx. pipiens* abundance + Potential avian species richness | 0.7288136 | 0.7272727 | 0.4560863 |

**Table S9.** Summary results of GAMs testing the effect of different mosquito biodiversity predictors on the probability of detecting WNV in mosquito pools. Models were fitted on the full dataset.

| Model | | | | | |
|---|---|---|---|---|---|
| **Mosquito species richness** | | **Estimate** | **Std. Error** | **z value** | **Pr(>\|z\|)** |
| | (Intercept) | 1.12619985 | 0.3392992 | 3.3191939 | 0.00090278 |
| | year2023 | -2.0692852 | 0.47667493 | -4.3410825 | 1.4178E-05 |
| | | **edf** | **Ref.df** | **Chi.sq** | **p-value** |
| | s(Sp_rich_mosq) | 1.759017407 | 2 | 9.546250942 | 0.004148832 |
| | s(Cropland) | 0.993454379 | 2 | 3.209652938 | 0.047145972 |
| | s(SparseVeg) | 1.369253657 | 2 | 3.603485468 | 0.069273941 |
| | s(Wetland) | 0.892193339 | 2 | 6.686504496 | 0.005883012 |
| | ti(Lon.Lat) | 3.8339E-05 | 16 | 8.84782E-06 | 0.958977632 |
| | | $R^2$ **(adj.)** = 0.350 | | **Deviance explained** = 31.8% | |
| **Mosquito evenness** | | **Estimate** | **Std. Error** | **z value** | **Pr(>\|z\|)** |
| | (Intercept) | 1.51669374 | 0.37643288 | 4.029121316 | 5.59857E-05 |
| | year2023 | -2.792366823 | 0.578256152 | -4.828944428 | 1.37259E-06 |
| | | **edf** | **Ref.df** | **Chi.sq** | **p-value** |
| | s(Evenness_mosq) | 0.916441457 | 2 | 9.649349951 | 0.000948007 |
| | s(Cropland) | 1.3604698 | 2 | 5.004236207 | 0.024667441 |
| | s(SparseVeg) | 0.782604175 | 2 | 3.249262471 | 0.039333438 |
| | s(Wetland) | 0.924639684 | 2 | 10.06184589 | 0.000847863 |
| | ti(Lon.Lat) | 0.000103044 | 16 | 5.34995E-05 | 0.730233859 |
| | | $R^2$ **(adj.)** = 0.355 | | **Deviance explained** = 31.5% | |
| ***Ae. albopictus* rel. abundance** | | **Estimate** | **Std. Error** | **z value** | **Pr(>\|z\|)** |
| | (Intercept) | 1.25656096 | 0.34909741 | 3.59945653 | 0.00031888 |

| | | | | | |
|---|---|---|---|---|---|
| | year2023 | -2.3117663 | 0.49026712 | -4.7153199 | 2.4133E-06 |
| | | **edf** | **Ref.df** | **Chi.sq** | **p-value** |
| | s(Prop_aealb) | 0.908633739 | 2 | 8.159713943 | 0.002548294 |
| | s(Cropland) | 0.657682983 | 2 | 1.796087747 | 0.097691699 |
| | s(SparseVeg) | 0.737331227 | 2 | 2.523794634 | 0.063749957 |
| | s(Wetland) | 0.844216844 | 2 | 4.513031504 | 0.020335584 |
| | ti(Lon.Lat) | 0.053175164 | 16 | 0.053268873 | 0.313063108 |
| | | **$R^2$ (adj.)** = 0.349 | | **Deviance explained** = 29.6% | |
| **Mosquito abundance** | | **Estimate** | **Std. Error** | **z value** | **Pr(>\|z\|)** |
| | (Intercept) | 1.50096493 | 0.3726527 | 4.02778495 | 5.6305E-05 |
| | year2023 | -2.7486874 | 0.5829855 | -4.714847 | 2.4189E-06 |
| | | **edf** | **Ref.df** | **Chi.sq** | **p-value** |
| | s(Ab_mosq) | 1.490111969 | 2 | 10.34626207 | 0.001146931 |
| | s(Cropland) | 0.000358604 | 2 | 0.000374997 | 0.305143862 |
| | s(SparseVeg) | 0.741369692 | 2 | 2.626512432 | 0.058064895 |
| | s(Wetland) | 0.714959225 | 2 | 2.067806442 | 0.086934477 |
| | ti(Lon.Lat) | 7.14869E-05 | 16 | 2.13368E-05 | 0.935581798 |
| | | **$R^2$ (adj.)** = 0.305 | | **Deviance explained** = 27.9% | |
| ***Cx. pipiens* rel. abundance** | | **Estimate** | **Std. Error** | **z value** | **Pr(>\|z\|)** |
| | (Intercept) | 1.269927813 | 0.342560958 | 3.707158638 | 0.000209598 |
| | year2023 | -2.303788067 | 0.506600661 | -4.547542561 | 5.42759E-06 |
| | | **edf** | **Ref.df** | **Chi.sq** | **p-value** |
| | s(Prop_cxpip) | 0.768512044 | 2 | 3.009509768 | 0.045502882 |
| | s(Cropland) | 1.095735198 | 2 | 4.557588829 | 0.021717808 |
| | s(SparseVeg) | 0.802636771 | 2 | 3.752280574 | 0.0287422 |

| | | | | | |
|---|---|---|---|---|---|
| | s(Wetland) | 0.900899789 | 2 | 7.465512013 | 0.003601358 |
| | ti(Lon.Lat) | 3.87707E-05 | 16 | 1.03086E-05 | 0.924064591 |
| | | **$R^2$ (adj.)** = 0.294 | | **Deviance explained** = 26.2% | |
| ***Cx. modestus*** **rel. abundance** | | **Estimate** | **Std. Error** | **z value** | **Pr(>\|z\|)** |
| | (Intercept) | 1.08617324 | 0.31829816 | 3.4124396 | 0.00064384 |
| | year2023 | -1.948511 | 0.44969913 | -4.3329214 | 1.4714E-05 |
| | | **edf** | **Ref.df** | **Chi.sq** | **p-value** |
| | s(Prop_cxmod) | 1.36068E-06 | 2 | 4.85802E-07 | 0.619747375 |
| | s(Cropland) | 0.812481298 | 2 | 4.100124809 | 0.023495357 |
| | s(SparseVeg) | 0.793244124 | 2 | 3.545863634 | 0.032958716 |
| | s(Wetland) | 0.879272536 | 2 | 6.066819243 | 0.008533006 |
| | ti(Lon.Lat) | 3.71886E-05 | 16 | 7.34171E-06 | 0.963345701 |
| | | **$R^2$ (adj.)** = 0.269 | | **Deviance explained** = 23.3% | |
| ***Oc. caspius*** **rel. abundance** | | **Estimate** | **Std. Error** | **z value** | **Pr(>\|z\|)** |
| | (Intercept) | 1.086173087 | 0.31829816 | 3.412439106 | 0.000643843 |
| | year2023 | -1.948510692 | 0.44969927 | -4.3329194 | 1.47145E-05 |
| | | **edf** | **Ref.df** | **Chi.sq** | **p-value** |
| | s(Prop_occas) | 4.84298E-06 | 2 | 1.06228E-06 | 0.748323508 |
| | s(Cropland) | 0.812482516 | 2 | 4.100121518 | 0.023495416 |
| | s(SparseVeg) | 0.793243948 | 2 | 3.545861839 | 0.032958761 |
| | s(Wetland) | 0.879271918 | 2 | 6.066811976 | 0.008533029 |
| | ti(Lon.Lat) | 3.16148E-05 | 16 | 5.86653E-06 | 0.962321641 |
| | | **$R^2$ (adj.)** = 0.269 | | **Deviance explained** = 23.3% | |

**Table S10.** Summary results of GAMs testing the effect of different mosquito biodiversity predictors on the probability of detecting WNV in mosquito pools. Models were fitted on the agricultural landscape dataset.

| Model | | | | | |
|---|---|---|---|---|---|
| **Mosquito species richness** | | **Estimate** | **Std. Error** | **z value** | **Pr(>\|z\|)** |
| | (Intercept) | 1.67563141 | 0.59365209 | 2.82258149 | 0.00476387 |
| | year2023 | -2.622224 | 0.76402465 | -3.4321196 | 0.00059888 |
| | | **edf** | **Ref.df** | **Chi.sq** | **p-value** |
| | s(Sp_rich_mosq) | 1.465533354 | 2 | 5.131906598 | 0.029011403 |
| | s(Imperv) | 1.54188286 | 2 | 5.096695723 | 0.029950436 |
| | s(SparseVeg) | 1.436866291 | 2 | 4.78786259 | 0.031008251 |
| | ti(Lon.Lat) | 0.735776515 | 15 | 2.438060464 | 0.057528435 |
| | | **$R^2$ (adj.)** = 0.449 | | **Deviance explained** = 43.4% | |
| **Mosquito evenness** | | **Estimate** | **Std. Error** | **z value** | **Pr(>\|z\|)** |
| | (Intercept) | 1.833931028 | 0.604157838 | 3.035516406 | 0.002401242 |
| | year2023 | -2.734653773 | 0.803249759 | -3.404487513 | 0.000662883 |
| | | **edf** | **Ref.df** | **Chi.sq** | **p-value** |
| | s(Evenness_mosq) | 1.531038874 | 2 | 4.049972624 | 0.068476914 |
| | s(Imperv) | 1.531289916 | 2 | 5.437456741 | 0.025585059 |
| | s(SparseVeg) | 0.850239399 | 2 | 2.732093625 | 0.05622018 |
| | ti(Lon.Lat) | 0.795469089 | 15 | 3.361979383 | 0.033982571 |
| | | **$R^2$ (adj.)** = 0.419 | | **Deviance explained** = 40.5% | |
| ***Ae. albopictus* rel. abundance** | | **Estimate** | **Std. Error** | **z value** | **Pr(>\|z\|)** |
| | (Intercept) | 1.86550186 | 0.62080185 | 3.00498763 | 0.00265592 |
| | year2023 | -2.732744 | 0.75714139 | -3.6092914 | 0.00030703 |
| | | **edf** | **Ref.df** | **Chi.sq** | **p-value** |

| | | | | | |
|---|---|---|---|---|---|
| | s(Prop_aealb) | 0.881013272 | 2 | 4.954598561 | 0.015092841 |
| | s(Imperv) | 1.043953662 | 2 | 2.07093964 | 0.125872689 |
| | s(SparseVeg) | 1.405079002 | 2 | 4.47255551 | 0.032000184 |
| | ti(Lon.Lat) | 1.408055331 | 16 | 3.670321624 | 0.058758362 |
| | | **$R^2$ (adj.)** = 0.435 | | **Deviance explained** = 42.3% | |
| **Mosquito abundance** | | **Estimate** | **Std. Error** | **z value** | **Pr(>\|z\|)** |
| | (Intercept) | 1.91046832 | 0.60532182 | 3.15612002 | 0.00159883 |
| | year2023 | -2.9432664 | 0.84966257 | -3.4640415 | 0.00053212 |
| | | **edf** | **Ref.df** | **Chi.sq** | **p-value** |
| | s(Ab_mosq) | 0.73530413 | 2 | 2.20654187 | 0.07571289 |
| | s(Imperv) | 1.41237986 | 2 | 3.55315995 | 0.07461789 |
| | s(SparseVeg) | 0.84046229 | 2 | 4.28709022 | 0.02085265 |
| | ti(Lon.Lat) | 0.80224471 | 16 | 3.55507435 | 0.02979496 |
| | | **$R^2$ (adj.)** = 0.358 | | **Deviance explained** = 35.7% | |
| ***Cx. pipiens* rel. abundance** | | **Estimate** | **Std. Error** | **z value** | **Pr(>\|z\|)** |
| | (Intercept) | 1.76336218 | 0.56936104 | 3.09708964 | 0.00195431 |
| | year2023 | -2.7343488 | 0.783065 | -3.4918542 | 0.00047968 |
| | | **edf** | **Ref.df** | **Chi.sq** | **p-value** |
| | s(Prop_cxpip) | 0.88775275 | 2 | 2.24799324 | 0.08910238 |
| | s(Imperv) | 1.64591926 | 2 | 7.4353279 | 0.00917152 |
| | s(SparseVeg) | 0.97940574 | 2 | 3.23160796 | 0.04470086 |
| | ti(Lon.Lat) | 0.76523746 | 16 | 2.82725097 | 0.04892402 |
| | | **$R^2$ (adj.)** = 0.377 | | **Deviance explained** = 37.3% | |
| ***Cx. modestus*** | | **Estimate** | **Std. Error** | **z value** | **Pr(>\|z\|)** |

| | | | | | |
|---|---|---|---|---|---|
| **rel. abundance** | (Intercept) | 1.610589012 | 0.544929314 | 2.955592536 | 0.003120691 |
| | year2023 | -2.363570739 | 0.701573294 | -3.368957683 | 0.00075453 |
| | | **edf** | **Ref.df** | **Chi.sq** | **p-value** |
| | s(Prop_cxmod) | 9.8153E-07 | 2 | 6.8453E-07 | 0.40382033 |
| | s(Imperv) | 1.6493981 | 2 | 7.13706983 | 0.01147708 |
| | s(SparseVeg) | 1.01530051 | 2 | 4.22280326 | 0.02463486 |
| | ti(Lon.Lat) | 0.78745906 | 16 | 3.34355899 | 0.03426021 |
| | | **$R^2$ (adj.)** = 0.337 | | **Deviance explained** = 33.7% | |
| ***Oc. caspius*** **rel. abundance** | | **Estimate** | **Std. Error** | **z value** | **Pr(>\|z\|)** |
| | (Intercept) | 1.491488104 | 0.540443434 | 2.759748773 | 0.005784583 |
| | year2023 | -2.190012328 | 0.712866082 | -3.072123059 | 0.002125421 |
| | | **edf** | **Ref.df** | **Chi.sq** | **p-value** |
| | s(Prop_occas) | 0.749738845 | 2 | 2.510574438 | 0.064782936 |
| | s(Imperv) | 1.602018567 | 2 | 6.276468938 | 0.017078593 |
| | s(SparseVeg) | 1.278259145 | 2 | 5.567882283 | 0.014046002 |
| | ti(Lon.Lat) | 0.748772375 | 16 | 2.565089812 | 0.057459974 |
| | | **$R^2$ (adj.)** = 0.385 | | **Deviance explained** = 38.2% | |

**Table S11.** Validation of GAMs testing the effect of different mosquito biodiversity predictors on the probability of detecting WNV in mosquito pools. Models were fitted both on the full dataset and on the agricultural landscape dataset.

| **Biodiversity variable** | **Dataset** | **TPR** | **TNR** | **TSS** |
|---|---|---|---|---|
| Mosquito species richness | Full dataset | 0.7457627 | 0.7090909 | 0.4548536 |
| | Agricultural landscape dataset | 0.6206897 | 0.6923077 | 0.3129973 |
| Mosquito evenness | Full dataset | 0.7118644 | 0.7272727 | 0.4391371 |
| | Agricultural landscape dataset | 0.6206897 | 0.6923077 | 0.3129973 |
| Mosquito abundance | Full dataset | 0.7288136 | 0.6727273 | 0.4015408 |
| | Agricultural landscape dataset | 0.5862069 | 0.5000000 | 0.0862069 |
| *Ae. albopictus* relative abundance | Full dataset | 0.7457627 | 0.7454545 | 0.4912173 |
| | Agricultural landscape dataset | 0.6206897 | 0.7307692 | 0.3514589 |
| *Cx. pipiens* relative abundance | Full dataset | 0.7288136 | 0.6363636 | 0.3651772 |
| | Agricultural landscape dataset | 0.7241379 | 0.6153846 | 0.3395225 |
| *Oc. caspius* relative abundance | Full dataset | 0.7627119 | 0.6363636 | 0.3990755 |
| | Agricultural landscape dataset | 0.7931034 | 0.6153846 | 0.4084881 |

**Table S12.** Summary results of GAMs testing the effect of both different avian biodiversity predictors and *Cx. pipiens* relative abundance on the probability of detecting WNV in mosquito pools. Models were fitted on the agricultural landscape dataset.

| Model | | | | | |
|---|---|---|---|---|---|
| ***Cx. pipiens* relative abundance + avian species richness** | | **Estimate** | **Std. Error** | **z value** | **Pr(>\|z\|)** |
| | (Intercept) | 2.073758607 | 0.688946538 | 3.010042857 | 0.002612108 |
| | year2023 | -3.487748814 | 0.973467998 | -3.582807878 | 0.000339921 |
| | points | -3.251088393 | 1.058517146 | -3.071361106 | 0.002130853 |
| | | **edf** | **Ref.df** | **Chi.sq** | **p-value** |
| | s(Prop_cxpip) | 3.24589E-06 | 2 | 5.81322E-07 | 0.810729111 |
| | s(Species_richness) | 1.737601688 | 2 | 10.84252051 | 0.002019141 |
| | s(Imperv) | 2.65947E-06 | 2 | 2.37095E-07 | 0.911454188 |
| | s(SparseVeg) | 0.797698365 | 2 | 3.309065163 | 0.038473855 |
| | ti(Lon.Lat) | 0.443837066 | 16 | 0.780321957 | 0.179926242 |
| | | **$R^2$ (adj.)** = 0.558 | | **Deviance explained** = 54.6% | |
| ***Cx. pipiens* relative abundance + avian functional richness** | | **Estimate** | **Std. Error** | **z value** | **Pr(>\|z\|)** |
| | (Intercept) | 1.580788741 | 0.54539127 | 2.898448924 | 0.003750134 |
| | year2023 | -2.581984318 | 0.759037891 | -3.401654053 | 0.000669794 |
| | points | -0.903806512 | 0.469756521 | -1.923989284 | 0.054355927 |
| | | **edf** | **Ref.df** | **Chi.sq** | **p-value** |
| | s(Prop_cxpip) | 0.42789747 | 2 | 0.527970653 | 0.275129465 |
| | s(Functional_richness) | 1.722405307 | 2 | 8.328648518 | 0.007189097 |
| | s(Imperv) | 3.54454E-06 | 2 | 1.90242E-06 | 0.50725975 |
| | s(SparseVeg) | 1.256627565 | 2 | 3.91031047 | 0.041674933 |
| | ti(Lon.Lat) | 0.618246734 | 16 | 1.527344408 | 0.110178631 |
| | | **$R^2$ (adj.)** = 0.397 | | **Deviance explained** = 39.4% | |

| | | | | | |
|---|---|---|---|---|---|
| ***Cx. pipiens* relative abundance**<br>+<br>**avian Shannon index** | | **Estimate** | **Std. Error** | **z value** | **Pr(>\|z\|)** |
| | (Intercept) | 2.405994108 | 0.73144863 | 3.289354864 | 0.001004173 |
| | year2023 | -3.324090022 | 0.926915139 | -3.586185919 | 0.00033555 |
| | points | -1.743046923 | 0.68283255 | -2.55267111 | 0.010690039 |
| | | **edf** | **Ref.df** | **Chi.sq** | **p-value** |
| | s(Prop_cxpip) | 1.44192E-05 | 2 | 7.42552E-06 | 0.580498877 |
| | s(Shannon) | 1.678992541 | 2 | 7.681101317 | 0.009618051 |
| | s(Imperv) | 1.521086641 | 2 | 5.547006192 | 0.020249271 |
| | s(SparseVeg) | 1.516272731 | 2 | 5.67051694 | 0.01647931 |
| | ti(Lon.Lat) | 1.483498846 | 16 | 5.041091092 | 0.016880992 |
| | | $R^2$ **(adj.)** = 0.493 | | **Deviance explained** = 50.8% | |
| ***Cx. pipiens* relative abundance**<br>+<br>**avian Simpson index** | | **Estimate** | **Std. Error** | **z value** | **Pr(>\|z\|)** |
| | (Intercept) | 1.871942645 | 0.593468924 | 3.154238693 | 0.001609174 |
| | year2023 | -2.796216824 | 0.800263 | -3.494122338 | 0.000475623 |
| | points | -0.353340666 | 0.409825411 | -0.862173638 | 0.388591974 |
| | | **edf** | **Ref.df** | **Chi.sq** | **p-value** |
| | s(Prop_cxpip) | 0.665549439 | 2 | 1.512418823 | 0.128564525 |
| | s(Simpson) | 0.723713041 | 2 | 1.653011617 | 0.128169469 |
| | s(Imperv) | 1.727915748 | 2 | 8.67282962 | 0.005179827 |
| | s(SparseVeg) | 1.232215226 | 2 | 3.775016736 | 0.044686136 |
| | ti(Lon.Lat) | 0.822240691 | 16 | 3.744715997 | 0.027709607 |
| | | $R^2$ **(adj.)** = 0.409 | | **Deviance explained** = 42.0% | |
| ***Cx. pipiens* relative abundance** | | **Estimate** | **Std. Error** | **z value** | **Pr(>\|z\|)** |
| | (Intercept) | 1.920371025 | 0.605096209 | 3.173662297 | 0.001505287 |

| | | | | | |
|---|---|---|---|---|---|
| + avian evenness | year2023 | -2.821996426 | 0.811159114 | -3.478967784 | 0.000503349 |
| | points | -0.232995648 | 0.383574729 | -0.607432218 | 0.543564113 |
| | | **edf** | **Ref.df** | **Chi.sq** | **p-value** |
| | s(Prop_cxpip) | 0.790409998 | 2 | 1.761576445 | 0.119751605 |
| | s(Evenness) | 0.706788079 | 2 | 1.679887737 | 0.117325037 |
| | s(Imperv) | 1.758437337 | 2 | 9.470022171 | 0.003322138 |
| | s(SparseVeg) | 1.198547643 | 2 | 4.322222561 | 0.028155801 |
| | ti(Lon.Lat) | 0.829901438 | 16 | 3.928592141 | 0.024532309 |
| | | **$R^2$ (adj.)** = 0.415 | | **Deviance explained** = 42.5% | |
| ***Cx. pipiens* relative abundance + potential avian species richness** | | **Estimate** | **Std. Error** | **z value** | **Pr(>\|z\|)** |
| | (Intercept) | 1.713848902 | 0.567750278 | 3.018666774 | 0.002538896 |
| | year2023 | -2.65293468 | 0.779939087 | -3.401463939 | 0.00067026 |
| | | **edf** | **Ref.df** | **Chi.sq** | **p-value** |
| | s(Prop_cxpip) | 1.037598126 | 2 | 2.135911634 | 0.121679035 |
| | s(pot.richness) | 0.453385046 | 2 | 0.75139685 | 0.186586613 |
| | s(Imperv) | 1.631415598 | 2 | 7.722280569 | 0.006745158 |
| | s(SparseVeg) | 0.792186975 | 2 | 3.141586306 | 0.041554505 |
| | ti(Lon.Lat) | 0.748448397 | 16 | 2.503648124 | 0.061237073 |
| | | **$R^2$ (adj.)** = 0.391 | | **Deviance explained** = 38.5% | |

**Table S13.** Validation of GAMs testing the effect of different avian biodiversity predictors combined with *Cx. pipiens* abundance on the probability of detecting WNV in mosquito pools. Models were fitted on the agricultural landscape dataset.

| **Biodiversity variable** | **TPR** | **TNR** | **TSS** |
|---|---|---|---|
| *Cx. pipiens* abundance + Avian species richness | 0.7931034 | 0.6923077 | 0.4854111 |
| *Cx. pipiens* abundance + Avian functional richness | 0.6896552 | 0.6153846 | 0.3050398 |
| *Cx. pipiens* abundance + Avian Shannon index | 0.6551724 | 0.6153846 | 0.2705570 |
| *Cx. pipiens* abundance + Avian Simpson index | 0.6896552 | 0.5769231 | 0.2665782 |
| *Cx. pipiens* abundance + Avian evenness | 0.6896552 | 0.6538462 | 0.3435013 |
| *Cx. pipiens* abundance + Potential avian species richness | 0.6896552 | 0.6153846 | 0.3050398 |
